# The crystal structure of bacteriophage λ RexA provides novel insights into the DNA binding properties of Rex-like phage exclusion proteins

**DOI:** 10.1101/2023.08.31.555690

**Authors:** Myfanwy C. Adams, Carl J. Schiltz, Jing Sun, Christopher J. Hosford, Virginia M. Johnson, Hao Pan, Peter P. Borbat, Jack H. Freed, Lynn C. Thomason, Carolyn Court, Donald L. Court, Joshua S. Chappie

## Abstract

RexA and RexB function as an exclusion system that prevents bacteriophage T4*rII* mutants from growing on *E. coli* λ phage lysogens. Recent data established that RexA is a non-specific DNA binding protein that can act independently of RexB to bias the λ bistable switch toward the lytic state, preventing conversion back to lysogeny. The molecular interactions underlying these activities are unknown, owing in part to a dearth of structural information. Here we present the 2.05-Å crystal structure of the λ RexA dimer, which reveals a two-domain architecture with unexpected structural homology to the recombination-associated protein RdgC. Modelling suggests that our structure adopts a closed conformation and would require significant domain rearrangements to facilitate DNA binding. Mutagenesis coupled with electromobility shift assays, limited proteolysis, and double electron-electron spin resonance spectroscopy support a DNA-dependent conformational change. I*n vivo* phenotypes of RexA mutants suggest that DNA binding is not a strict requirement for phage exclusion but may directly contribute to modulation of the bistable switch. We further demonstrate that RexA homologs from other temperate phages also dimerize and bind DNA *in vitro*. Collectively, these findings advance our mechanistic understanding of Rex functions and provide new evolutionary insights into different aspects of phage biology.

## INTRODUCTION

Bacteriophage λ has served as a powerful experimental system since its discovery by Esther Lederberg (1) and has yielded ground-breaking insights into many fundamental processes in molecular biology (2). As a temperate phage, λ exhibits a bi-modal life cycle of infection where the viral chromosome can either stably integrate into the host *Escherichia coli* chromosome to form a lysogen and replicate passively as a prophage, or the phage can switch to a lytic state wherein viral gene expression commandeers the bacterial cellular machinery to produce and assemble virions, ultimately lysing and killing the host as a means of escape (3). Following integration and establishment of the λ prophage, lysogeny is maintained by the phage CI repressor, which binds two control regions (OL and OR) to repress transcription from the leftward and rightward lytic promoters P_L_ and P_R_ while concomitantly directing its own expression from the maintenance promoter P_RM_ (4, 5) (**Supplementary Figure S1**). Each OL and OR control region contains three operator sites: OL1, OL2, OL3 and OR1, OR2, OR3 respectively. CI repressor dimers preferentially bind OL1 and OR1 with high affinity and direct the cooperative binding of a second dimer to OL2 / OR2 via stabilizing interactions in the repressor C-terminal domain (6). Inherent structural asymmetry precludes the binding of a third CI dimer at the adjacent OL3 and OR3 sites (7). Alternative cooperative binding of CI dimers to OR2 and OR3 can occur if the OR1 operator site is mutated (8). Stronger repression occurs through long range DNA looping and the formation of CI octamers that tether OL1 and OL2 to OR1 to OR2 (9–12) (**Supplementary Figure S1A**), which can also facilitate cooperative binding of CI dimers to OL3 and OR3 at lower occupancy in an alternative tetrameric arrangement (**Supplementary Figure S1A**). Prophage induction and transition to the lytic state can be triggered by cellular insults such as DNA damage that induce the host cell’s SOS response (**Supplementary Figure S1B**). RecA-dependent proteolytic CI auto-cleavage during SOS reverses prophage repression and vacates the operator sites, enabling transcription from the P_L_ and P_R_ lytic promoters (13–17). The first gene in the P_R_ operon, *cro*, encodes the Cro repressor, which binds to the same six operator sites as the CI repressor, but with a different affinity pattern (18, 19). Cro dimers have the highest affinity for OR3 and once bound, repress transcription from the P_RM_ promoter and prevent further CI expression, stabilizing the switch to lytic growth and locking in lytic gene expression (20, 21).

The *rex* genes are situated directly adjacent the *cI* repressor gene in the λ genome and are transcribed from the same P_RM_ maintenance promoter that drives constitutive expression of the repressor during lysogeny (**Supplementary Figure S1A**). They produce a 31 kDa soluble protein, RexA, and a 16 kDa protein, RexB. RexB has a predicted four transmembrane helical topology and localizes to the bacterial inner membrane (22). RexA and RexB physically interact *in vivo* (23, 24) and together inhibit productive replication of bacteriophage T4 *rII* mutants (21–24) in λ lysogens (25–27). This observation has resulted in the proposal that together RexA and RexB form an anti-phage defense system (22, 25, 28). The Rex proteins additionally associate with CI and Cro and influence λ phage development (23, 24). We previously showed that RexA is a non-specific DNA binding protein that stabilizes the CI-Cro bistable switch in the lytic position, thereby helping prevent conversion back to lysogeny (24) (**Supplementary Figure S1C**). This RexA-mediated stabilization of lytic growth occurs independently of RexB but can be further modulated when RexB is present. Co-expression of RexA and RexB can also cause energetic defects in host cells (22, 29) and both proteins interact with subunits of the *E. coli* NADH dehydrogenase when analyzed using the bacterial adenylate cyclase two-hybrid system (BACTH) (30). Genetic and biochemical studies have demonstrated other independent functions for each protein: RexB can inhibit ATP-dependent ClpX and ClpA proteolysis of the λ replication protein O, the antitoxin proteins Phd from plasmid prophage P1, and MazE from *E. coli* (31–34) while RexA interacts with the *E. coli* sulfur metabolism protein CysN in BACTH assays (30). Although a cogent model connecting various Rex effects to inhibition of host energetics has been proposed (30), detailed molecular mechanisms for many functions remain a mystery, owing in part to a lack of structural information.

Here we present the crystal structure of the λ RexA dimer at 2.05-Å resolution. Each RexA monomer contains a split globular domain and a dimerization domain, which display unexpected structural homology to the bacterial recombination-associated protein RdgC. Structural superposition and modelling suggest that we captured RexA in a closed configuration and that a significant reorganization of the globular domains would be required for DNA binding. Mutagenesis coupled with electromobility shift assays, limited proteolysis, and double electron-electron spin resonance spectroscopy (DEER) support the possibility of DNA-dependent conformational change. We observe no correlation between RexA’s ability to bind DNA *in vitro* and phage exclusion activity *in vivo*; however, the RexA D215W conformational mutant that exhibits enhanced DNA binding can promote transition to the lytic state in a genetic background where lysogeny normally prevails. We further demonstrate that RexA homologs from other temperate phages also dimerize and can bind DNA *in vitro*. Collectively, these data advance our mechanistic understanding of Rex functions and provide new evolutionary insights into the regulation of lysogenic-lytic transitions and anti-phage defense activities encoded in the genomes of temperate bacteriophages.

## MATERIAL AND METHODS

### Cloning, expression, and purification of bacteriophage lambda RexA

DNA encoding bacteriophage λ RexA protein (UniProt P68924) was codon optimized for *E. coli* expression and synthesized commercially by Bio Basic Inc. DNA encoding full-length RexA (residues 1-279) was amplified by PCR and cloned into pET21b, introducing a 6xHis tag at the C-terminus. Native RexA was transformed into BL21(DE3) cells, grown at 37°C in Terrific Broth to an OD_600_ of 0.8-1.1, and then induced with 0.3 mM IPTG overnight at 19°C. Cells were pelleted, washed with nickel loading buffer (20 mM HEPES pH 7.5, 500 mM NaCl, 30 mM imidazole, 5% glycerol (v:v), and 5 mM β-mercaptoethanol), and pelleted a second time. Pellets were frozen in liquid nitrogen and stored at -80°C. Selenomethionine labeled (SeMet) RexA was expressed in minimal media in absence of auxotrophs as described previously (35).

Thawed 500 ml pellets of native and SeMet RexA constructs were resuspended in 30 ml of nickel loading buffer supplemented with 5 mg DNAse, 5 mM MgCl_2_, 10 mM PMSF, and a Roche complete protease inhibitor cocktail tablet. Lysozyme was added to a concentration of 1 mg/ml and the mixture was incubated for 10 minutes rocking at 4°C. Cells were disrupted by sonication and the lysate was cleared via centrifugation at 13 000 rpm (19 685 g) for 30 minutes at 4°C. The supernatant was filtered, loaded onto a 5 ml HiTrap chelating column charged with NiSO_4_, and then washed with nickel loading buffer. Native and SeMet RexA were eluted by an imidazole gradient from 30 mM to 1 M. Pooled fractions were dialyzed overnight at 4°C into S loading buffer (20 mM HEPES pH 7.5, 50 mM NaCl, 1 mM EDTA, 5% glycerol (v:v), and 1 mM DTT). Significant precipitation occurred during dialysis; however, RexA was never observed in the insoluble fraction. The dialyzed sample was applied to 5 ml HiTrap SP column equilibrated with S loading buffer, washed in the same buffer, and eluted with a NaCl gradient from 50 mM to 1 M. Peak fractions were pooled, concentrated, and further purified by size exclusion chromatography (SEC) using a Superdex 75 16/600 pg column (Cytiva). Native and SeMet RexA were exchanged into a final buffer of 20 mM HEPES pH 7.5, 150 mM KCl, 5 mM MgCl_2_, and 1 mM DTT during SEC and concentrated to 10-70 mg/ml. All mutations were introduced via QuikChange site directed mutagenesis using Pfu Turbo polymerase with exact primer overlap. Mutant RexA proteins were purified as described for the wildtype native protein.

### Cloning, expression, and purification of RexA homologs: Sbash gp30, CarolAnn gp44, and Toast gp42

RexA homologs were identified through BLAST against the Actinobacteriophage Database (PhagesDB) (36), the Integrated Microbial Genomes database (37), and the KEGG database (38). DNA encoding RexA homologs from mycophages Sbash (*gp30*), CarollAnn (*gp44*), and Toast (*gp42*) were codon optimized for *E. coli* expression and synthesized commercially by Twist Biosciences. Full-length constructs for each protein (Sbash 30: residues 1-372; CarolAnn 44: residues 1-370; Toast 42: residues 1-337) were inserted via Gibson cloning into pET15bP. This vector introduced an N-terminal 6x-His tag, cleavable by protease HRV 3C. Constructs were transformed into BL21(DE3) cells, grown at 37°C in Terrific Broth to an OD_600_ of 0.8-1.1, and then induced with 0.3 mM IPTG overnight at 19°C. Cells were pelleted and resuspended in nickel loading buffer (20 mM HEPES pH 7.5, 500 mM NaCl, 30 mM imidazole, 5% glycerol (v:v), and 5 mM β-mercaptoethanol) supplemented with 0.5 mg DNAse, 10 mM MgCl_2_, 1 mM PMSF, and a Roche complete protease inhibitor cocktail tablet. Lysozyme was added to a concentration of 1 mg/ml and the mixture was incubated for 10 minutes rocking at 4°C. Cells were disrupted by sonication and the lysate was cleared via centrifugation at 13 000 rpm (19 685 g) for 30 minutes at 4°C. The supernatant was filtered, loaded onto a 5 mL HiTrap chelating column charged with NiSO4, and then washed with nickel loading buffer. RexA homologs were eluted by an imidazole gradient from 30 mM to 500 mM. HRV 3C protease was added to the pooled fractions, which were subsequently dialyzed overnight at 4°C against Q buffer (20 mM Tris-HCl pH 8.0, 50 mM NaCl, 1 mM EDTA, 5% glycerol (v:v), and 1 mM DTT). The clarified protein sample was applied to a 5 ml HiTrap Q column equilibrated with Q buffer. The sample was washed with Q buffer and eluted along a NaCl gradient from 50 mM to 500 mM over 10 column volumes. Peak Q column fractions were pooled, concentrated, and further purified by SEC using a Superdex 200 10/300 GL sizing column equilibrated in 20 mM HEPES pH 7.5, 150 mM KCl, and 1 mM DTT. Peak SEC fractions were concentrated to 10-20 mg/ml, flash frozen in liquid nitrogen, and stored at -80°C.

### Size exclusion chromatography coupled to multi-angle light scattering (SEC-MALS)

Purified RexA constructs and homologs at 4 mg/mL were subjected to size-exclusion chromatography using a Superdex 200 10/300 GL Increase column (Cytiva) equilibrated in SEC buffer (20 mM HEPES pH 7.5, 150 mM KCl, and 1mM DTT). The column was coupled to a static 18-angle light scattering detector (DAWN HELEOS-II) and a refractive index detector (Optilab T-rEX) (Wyatt Technology). Data were collected continuously at a flow rate of 0.6 mL/min. Data analysis was carried out using the program Astra VI and graphs generated using Kaleidagraph (Synergy Software). Monomeric BSA at 5 mg/mL (Sigma) was used for normalization of the light scattering detectors and data quality control.

### Crystallization, X-ray data collection, and structure determination

SeMet RexA at 12 mg/ml was crystallized by sitting drop vapour diffusion at 20°C in 1.9 M AmS04 and 225 mM NDSB-195 (Hampton Research). Crystals were of the space group P32 2 1 with unit cell dimensions a = 56.4 Å, b = 56.4 Å, c = 326.22 Å and α = 90°, β = 90°, γ = 120° and contained a dimer in the asymmetric unit. Samples were cryoprotected by transferring the crystal directly to 25% sucrose prior to freezing in liquid nitrogen. Crystals were screened and optimized at the MacCHESS F1 beamline at Cornell University and single-wavelength anomalous diffraction (SAD) data (39) were collected remotely on the tuneable NE-CAT 24-ID-C beamline at the Advanced Photon Source at the selenium edge (λ=0.9792Å) at 100K to a resolution of 2.68 Å (**Supplementary Table S1**). Data were integrated and scaled via the NE-CAT RAPD pipeline, using XDS (40) and AIMLESS (41), respectively. A total of 14 selenium sites – seven per monomer – were found using SHELX (42) and used for initial phasing. Density modification and initial model building was carried out using the Autobuild routines of the PHENIX package (43). Further model building and refinement was carried out manually in COOT (44) and PHENIX (43). Xtriage (45) analysis indicated that SeMet RexA crystals were twinned with an estimated twin fraction of 0.140. The twin law -h,-k,l was thus applied during all refinement procedures. The resulting model was incomplete and lacking interpretable density for residues 32-41, 60-79, 192-196, and 227-246. A second crystal form was obtained by sitting drop vapor diffusion at 20°C in 2.2 M AmS0_4_ and 0.2M CdCl_2_. These crystals were cryoprotected by placing 1:1 v/v 20% sucrose on drop and then transferring to 30% sucrose prior to freezing. Crystals were of the space group P32 2 1 with similar unit cell dimensions a = 55.65 Å, b = 55.65 Å, c = 322.12 Å and α = 90°, β = 90°, γ = 120° and similarly contained a dimer in the asymmetric unit. Diffraction data were collected on the NE-CAT 24-ID-C beamline at the Advanced Photon Source at the selenium edge (λ =0.9792Å) at 100K and integrated and scaled as described above (**Supplementary Table S1**). These crystals showed improved diffraction though were similarly twinned. The twin fraction of 0.090 and the twin law -h,-k,l was similarly applied. Crystal form 2 was solved by molecular replacement with PHASER (46) using the SAD derived structure from crystal form 1 as the search model. Crystal form 2 was refined to 2.05Å resolution with R_work_/R_free_ values of 20.63 % / 25.12 % (**Supplementary Table S1**). The final model contained residues 1-30 and 34-279 in chainA, residues 1-279 in chain B, 364 water molecules, eight sulfates, and nine cadmium ions.

Structural superpositions and interpolations for molecular morphing were carried out in Chimera (47), while surface electrostatics were calculated using APBS (48). All structural models were rendered using Pymol (Schrodinger). Visual mapping of conserved residues was carried out using the ConSurf server (49).

### Preparation of oligonucleotide substrates

All DNA oligonucleotides were synthesized commercially by Integrated DNA Technologies (IDT). Lyophilized single-stranded oligonucleotides were resuspended to 200 µM in 20 mM HEPES pH 7.5 and stored at -20°C. For EMSA visualisation the upper strand oligonucleotides included a 6-FAM modification on the 5’ end. Duplex substrates were prepared by heating equimolar concentrations of complementary strands (denoted as ‘us’ and ‘ls’ indicating upper and lower strands) to 95°C for 5 minutes followed by cooling to room temperature overnight and then purification on an S-300 spin column (GE) to remove single stranded DNA. Sequences for all substrates can be found in **Supplementary Table S2**.

### Electrophoretic mobility shift assays (EMSA)

Binding was performed using either PCR generated or commercially synthesized DNA substrates (**Supplementary Table S2**) with purified protein. Increasing concentrations of protein (0 – 12.5 / 25 □M) were incubated with 500 nM DNA. Samples were prepared in EMSA buffer (20 mM Tris-HCl pH 8.0, 50 mM NaCl) to a total reaction volume of 20 µL and incubated at room temperature for 30 minutes. RexA EMSAs were run with 5’-6FAM oligos on a hand poured 10% 20 cm x 20 cm polyacrylamide gel. These gels were run at 4°C on a large format PROTEAN II XL Cell system for 4 hours at 120V in 1X TAE (40 mM Tris, 40 mM acetate, 1 mM EDTA, pH 8.0) buffer and imaged on a ChemiDoc imager using the “fluorescein” setting. CarolAnn, Toast and Sbash EMSAs used the same reaction conditions as above but with unlabelled oligos, loaded onto a 12% precast polyacrylamide gel (BioRad), and run at 100V for 90 minutes in 1X TAE buffer at room temperature. Gels were stained with SYBR Gold (ThermoFischer) in 1X TAE for 20 min at room temperature and visualized using a Bio-Rad Gel Doc EZ imager system.

### Limited proteolysis

Assays were carried out in proteolysis buffer (50 mM NaCl and 20 mM Tris-HCl pH 8.0) with a final volume of 15 µL. Proteins (final concentration 1 mg/mL) and DNA (final concentration 25 µM) were first allowed to incubate at room temperature 30 minutes before enzyme was added to a final concentration of 50 µg/mL. Samples were incubated at room temperature for an additional 30 minutes before being run on a polyacrylamide gradient gel (4-20%, Novex) at 150V for 45 minutes.

### Double electron-electron resonance (DEER) spectroscopy

100 µM of protein was incubated with MTSL (*S*-(1-oxyl-2,2,5,5-tetramethyl-2,5-dihydro-1H-pyrrol-3-yl)methyl methanesulfonothioate, Toronto Research Chemicals) nitroxide spin label to a final 1:10 protein-to-label molar ratio, diluted in SEC (20 mM HEPES pH 7.5, 150 mM KCl) buffer, and incubated overnight at 4°C. Unreacted label was removed by washing the protein several times in a concentrator, labelled protein was adjusted to 100 µM final concentration in deuterated SEC buffer lacking DTT before 8 mg of Gly-D8 was added. 20 μL spin-labeled samples were loaded into 2 mm I.D. quartz capillaries (Vitrocom) and snap frozen in liquid nitrogen prior to DEER measurements. DEER measurements were performed at 60 K using a home-built Ku band 17.3 GHz pulse ESR (50, 51). A four-pulse DEER sequence (52) was used routinely with the detection and pump π-pulses having respective widths of 32 ns and 16 ns. The detection pulses were applied at the low-field spectral edge, pumping was at the central maximum. A 32-step phase cycle (53) suppressed unwanted contributions to the signal. Nuclear modulation effects from surrounding protons were suppressed by summing 4 data traces recorded with inter-pulse separations incremented by 9.5 ns in subsequent measurements (54). Time-domain DEER data were reconstructed into distance distributions using standard approaches for baseline removal (52, 55, 56). The distances between spin pairs were reconstructed with L-curve Tikhonov regularization (57). Continuous wave (CW-ESR) spectra were recorded using a Bruker E500 X-band spectrometer at 200 K, modulation amplitude 0.2 mT, mw power 0.6 mW.

### Generation and culture of bacterial strains and phage stocks

Bacterial strains (**Supplementary Table S3**) were constructed using a combination of recombineering and P1 transduction methods (58, 59). Sequences of single-strand oligonucleotides used for recombineering are available upon request. RexA mutations (R219A/K221A, Δ239-244, and D215W) were introduced into a previously established P_L_P_R_ dual reporter that was constructed by the insertion of the λimmunity region into the *E. coli lac* operon (21). Reporter strains contain the temperature-sensitive *cI857* repressor with the P_R_ lytic promoter driving expression *lacZ* and the P_L_ lytic promoter driving expression of the firefly luciferase gene. In a control strain, the *rex* genes were replaced with a chloramphenicol resistance cassette, and this strain was also assayed, along with a wildtype *E. coli* MG1655 that lacks the reporter system.

Bacterial cultures were grown in L broth containing 10 g tryptone, 5 g yeast extract and 5 g NaCl per liter, and on L plates, which contained ingredients above and 1.5% Difco agar. Cultures for plating phage were grown to exponential phase in tryptone broth containing 10 g tryptone, 5 g NaCl, and 10mM MgSO_4_ per liter. Phage stocks were maintained in TMG, containing 10 mM Tris base, 10 mM MgSO_4_ and 0.01% gelatin at pH 7.4. Phage were enumerated on tryptone plates 10 g tryptone and 5 g NaCl per liter using 0.25 ml of fresh plating cultures mixed with 2.5 ml melted tryptone top agar (0.7% agar) containing 10 g tryptone and 5 g NaCl. MacConkey Lactose agar medium was from Difco and contained 1% lactose and 1.35% agar. Dilutions of bacteria were made in M9 Salts while dilutions of phage were made in TMG.

### Phage exclusion assays

High titer stocks of bacteriophages λ, T4, and T4*r*II were serially diluted in 10-fold increments into TMG. A 10 μl aliquot of each phage dilution was spotted on tryptone petri plates bearing freshly hardened lawns of top agar containing freshly grown cultures of the appropriate bacterial strains. Petri plates were allowed to air dry with the lids off. Once the phage spots dried, petri plates were inverted and incubated at 32°C overnight.

### Papillation assays

The papillation of *E. coli* strains carrying the dual P_L_ P_R_ reporters with a *c*I*857 ind1* allele and *rexA* mutations was examined by plating dilutions of fresh LB overnight cultures on MacConkey-Lactose to obtain isolated single colonies. Both Cro^+^ and *cro27* versions of the reporters were examined. MacConkey-Lactose plates were incubated for several days at 32-34°C until papillae arose within individual colonies. All colonies are white after one day of incubation but develop red papillae after two to three days. In the Cro^+^ strains, these papillae arise by an epigenetic transition to the lytic state in individual cells within the colony and consequent Cro-dependent expression of *lacZ* from the P_R_ promoter. In *cro27* strains, the low numbers of red papillae arise due to mutations in the *c*I repressor gene (24). The plates were photographed and the number of papillae in individual colonies was counted for each reporter strain (see **Supplementary Table S3** for genotypes). At least one hundred colonies were scored for each genotype and the data were plotted as scatterplots using GraphPad Prism, with each small vertical line indicating the number of papillae found in a single colony. Mean and standard deviation were calculated automatically in Prism. Numerical data and statistics can be found in the associated raw data file.

## RESULTS

### Structure of bacteriophage λ RexA reveals a two-domain fold

The RexA dimer readily crystallized in two unique crystal forms. Crystal form 1 was initially solved to 2.68-Å resolution by SAD phasing (39) using selenomethionine-labeled protein (35). The resulting structure, however, was incomplete and was thus used as a search model for molecular replacement to solve crystal form 2, which refined to 2.05-Å resolution with an R_work_/R_free_ of 20.6%/25.1%. The final model shows a symmetric dimer that measures 95 Å along the longest axis. Each monomer contains a split globular domain (residues 1-139 and 248-279) comprised of a seven-stranded antiparallel β-sheet flanked by α-helices that is divided by a dimerization domain consisting of a four-stranded antiparallel β-sheet with four α-helices localized on one face (residues 143-234) (**Fig. 1A,B**). These domains are connected via two flexible linkers that we term the hinge loop (140–142) and the swivel loop (235–247) (**Fig. 1B,C**). The β-sheets of the dimerization domain pack together to form a continuous eight-stranded anti-parallel platform above which helices α6, α7, and η1 interdigitate (**Fig. 1D**). These segments are primarily stabilized through extensive hydrophobic interactions (**Fig. 1E**) and a set of hydrogen bonds along the length of the β10-β10 interstrand interface (**Fig. 1F**), which together yield a total buried surface area of 3059Å^2^. A few additional hydrogen bonds are scattered across the dimer interface, including interactions between T176 and D204, D187 and K191, R196 and the backbone carbonyl of E181, and N208 and the backbone carbonyl of L173 (**Fig. 1F**). The remaining polar residues within the dimerization domain are primarily oriented outward into solution and interact with water molecules on the protein surface.

**Fig. 1.**
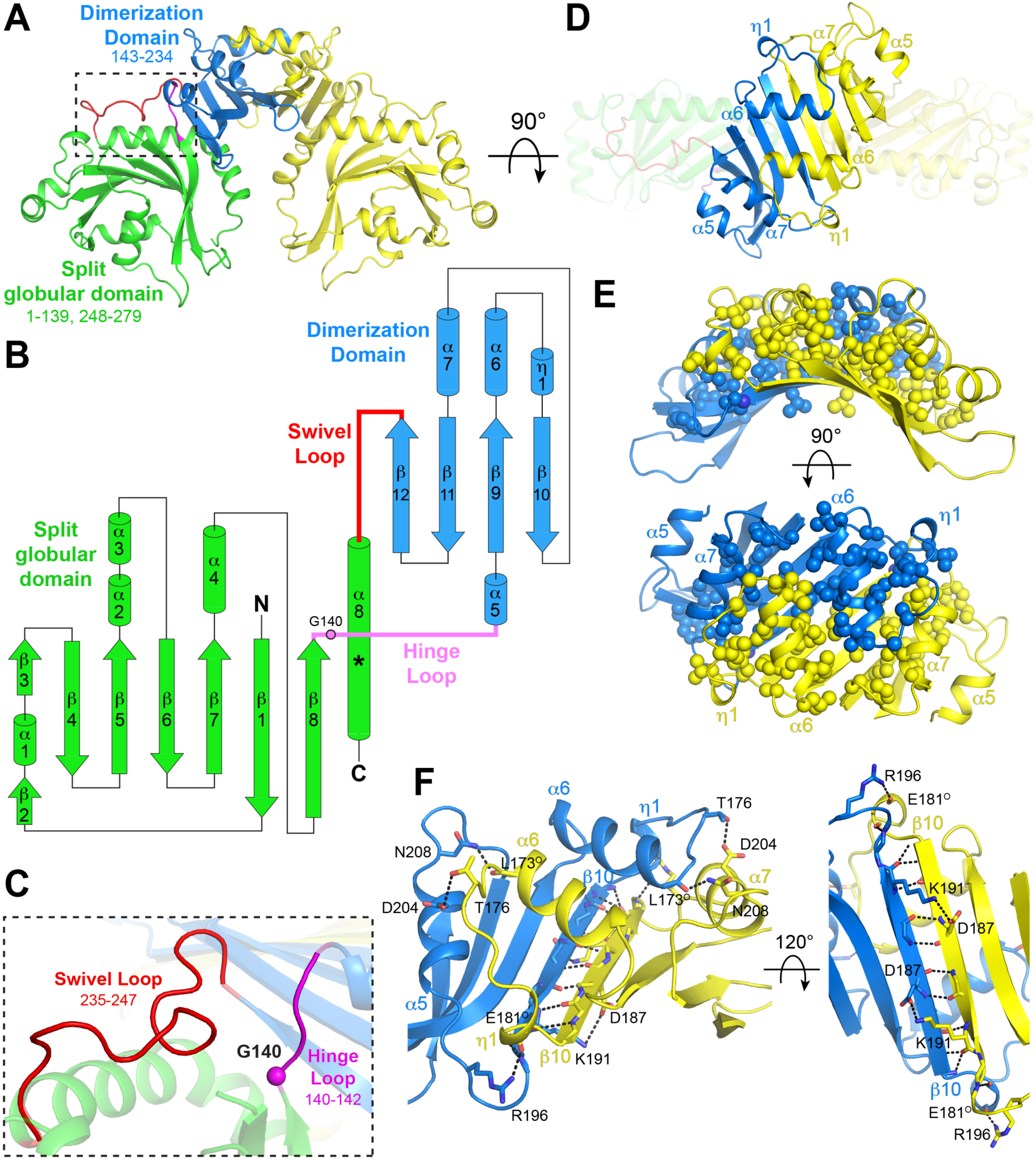
Structure of RexA. A. Structure of λ RexA dimer. Split globular domain and dimerization domain are labeled in one monomer are colored green and blue respectively. Second monomer is colored yellow for contrast. Dashed box highlights flexible loop connections (pink and red) between domains (see panel C). B. Topology diagram of RexA monomer. Pink circle indicates putative G140 hinge and asterisk denotes position of the kink present in the C-terminal α9 helix. C. Zoomed view of box in A showing conformations of the hinge loop (pink) and swivel loop (red). Pink sphere shows position of putative G140 hinge. D. Dimerization interface of RexA. Interdigitating helices are labeled. E, F. Hydrophobic (E) and hydrogen bonding (F) interactions stabilize the RexA dimer interface. Hydrophobic side chains are shown as spheres and hydrogen bonds are depicted as dashed black lines. Participating side chains are labeled, with “O” superscript denoting backbone carbonyl oxygen.

The globular domains are situated below the dimerization domains and have only minimal contact across the 2-fold symmetry axis via the C-terminus of the α8 helix. This helix kinks at residues 261-263 and interacts with strand β11 of its affiliated dimerization domain, anchored by a salt bridge between D215 and R61 (**Supplementary Fig. S2A**). A bound sulfate and two cadmium ions from the crystallization solution provide further hydrogen bonding contacts in this region in monomer A (**Supplementary Fig. S3A and 3B, panels I and II**). Additional cadmium and sulfate ions associate with the surface of the RexA dimer (**Supplementary Fig. S3A**), either binding to various pockets within each monomer (**Supplementary Fig. S3B**) or forming intermolecular ionic interactions that mediate crystal lattice contacts (**Supplementary** Fig. 3C). The sandwiching of a cadmium ion between two monomers at the two-fold symmetry axis (**Supplementary Fig. S3C, panel I**) produced a significant change in packing relative to crystal form 1, which ultimately yielded the high-resolution diffraction.

### Structural homology with RdgC implicates a conserved mode of DNA binding

Proteins with similar activities often evolve from a common ancestor and share a conserved structural fold. Fold matching and structural alignment are thus useful tools for deducing functional properties and identifying important structural motifs in poorly characterized proteins (60). Initial sequence-based fold prediction with Phyre2 (61) and I-TASSER (62) failed to identify reliable structural homologs, likely owing to the divided nature of the globular domain. Using the DALI alignment algorithm (63), we identified the bacterial recombination-associated protein RdgC as a structural relative of RexA (Z score 4.4, RMSD 2.7). *E. coli* RdgC reduces RecA-catalyzed strand exchange, RecA ATPase activity, and RecA-dependent cleavage of the SOS transcriptional repressor LexA (64). Deletion of *rdgC* is toxic in Δ*priA* and Δ*priB* strains, suggesting it also plays an important role in PriA/PriB-dependent replication restart following repair and processing of DNA damage (65, 66). RdgC homologs form ring shaped dimers (67–69) that bind DNA non-specifically (65, 68, 70). Each RdgC monomer contains a base domain consisting of a five-stranded anti-parallel β-sheet flanked by four α-helices, a globular center domain, and a tip domain comprised of two α-helices spliced between the β4 and β5 strands of the center domain (**Supplementary Fig. S4A,B**). The base and tip domains mediate dimerization (67–69), acting as contact points that hold the ring together (**Supplementary Fig. S4C**). RexA’s split globular domain and dimerization domain structurally align with the RdgC central and base domains respectively (**Fig. 2A,B**), each sharing a similar topology (**Fig. 1B, Supplementary Fig. S4B**).

**Fig. 2.**
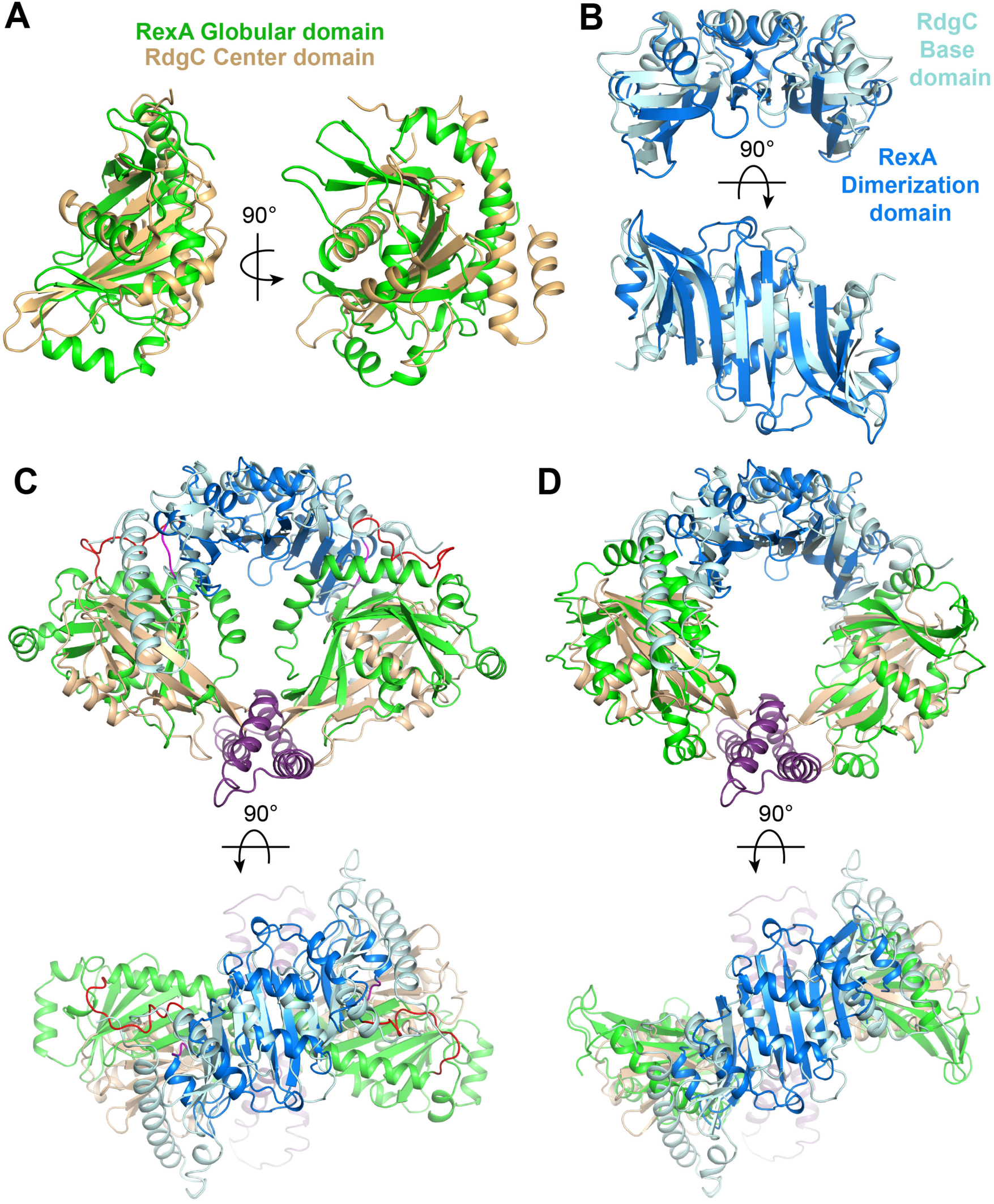
RexA shares structural homology with RdgC. A. Superposition of RexA split globular domain (green) and RdgC center domain (beige). B. Superposition of RexA dimerization domains (blue) and RdgC base domains (pink). C. Superposition of crystallized RexA dimer with crystallized RdgC dimer. Structures aligned via the dimerization/base domains and individual domains are colored as in A and B. D. Structural alignment of individual RexA domains onto RdgC dimer yields a modeled ‘open’ conformation.

The RexA dimer contains an extensive basic patch that stretches across the surface of the globular domains at the dimer interface (**Fig. 3A**). How this might be utilized to bind negatively charged DNA remains unclear. Previous studies suggest that RdgC binds DNA in the central pore of its ring-shaped dimer (67, 69). Calculation of surface electrostatics shows that this pore in RdgC is lined with basic residues that can interact with the negatively charged DNA backbone (**Fig. 3D**). Superposition of RexA and RdgC shows that the orientation of the globular domains in the crystallized RexA dimer occludes the space beneath the dimerization interface, blocking the formation of an analogous pore and preventing DNA from associating in the same manner (**Fig. 2C, 3E and 3H**). From the observed homology between the individual domains (**Fig. 2A,B**), however, we can model an open conformation for RexA that adopts a horseshoe shape with a large open cavity (**Fig. 2D, 3B and 3F**). The resulting arrangement of RexA domains places basic side chains along the inner surface of the open cavity (**Fig 3B**). Consurf analysis (49) reveals that the most highly conserved residues among putative RexA sequences also line this cavity in the modeled open state while poorly conserved side chains localize on the exterior of the structure (**Fig. 3I,J and Supplementary Fig. S5**). Strikingly, an unbiased model of the RexA dimer generated using AlphaFold-Multimer (71) predicts a similar open conformation (**Fig. 3G**) with comparable spatial arrangement of both surface charges (**Fig. 3C**) and conserved residues (**Fig. 3K**). Based on this modeling, we predict RexA undergoes a major conformational change that reorients the globular domains to facilitate DNA binding.

**Fig. 3.**
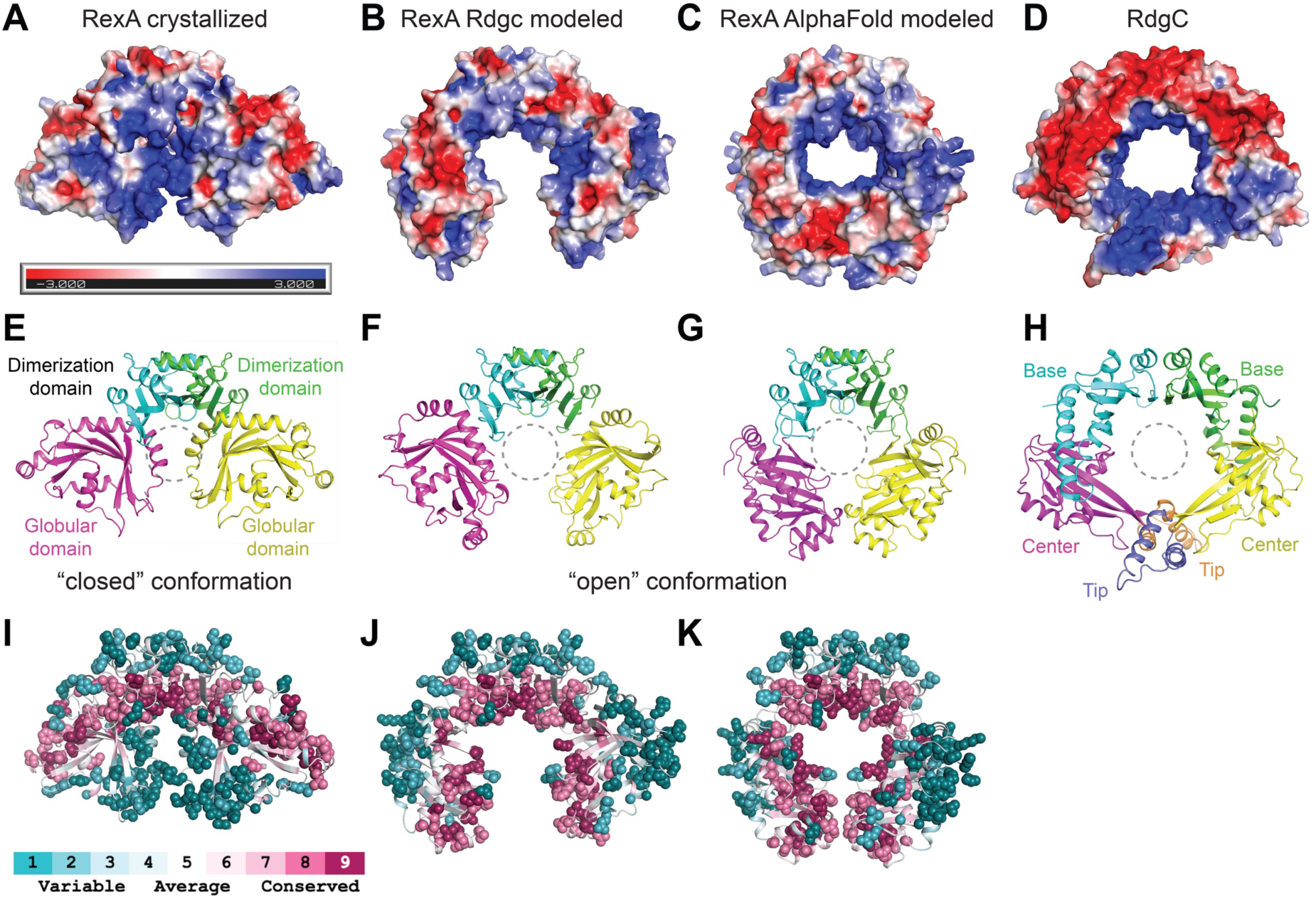
Putative RexA conformational change suggested by RdgC and AlphaFold modeling. A-D. Electrostatic surfaces of (left to right) crystallized RexA dimer (A), modeled open RexA dimer based on RdgC homology (B), modeled RexA dimer predicted via AlphaFold-Multimer (71) (C), and RdgC (PDB: 2OWL) (D). Scale bar indicates electrostatic surface coloring from -3 K_b_T/e_c_ to +3 K_b_T/e_c_. E-H. Domain organization of crystallized RexA dimer (E), modeled open RexA dimer based on RdgC homology (F), modeled RexA dimer predicted from AlphaFold-Multimer (71) (G), and RdgC (H). Structurally analogous domains are similarly colored to highlight their relative positions in each structure. Dashed circle shows the relative position of the RdgC central pore in each structure based on superposition (see Fig. 2C**,D**) through which DNA is thought to pass (69). I-K. Distribution of conserved residues on the crystallized (I), RdgC-modeled (J), and AlphaFold-modeled (K) RexA dimer structures. Coloring generated using the ConSurf server (49) and the sequence alignment in **Supplementary Fig. S4**.

### Mutational analysis supports a DNA-dependent conformational change in RexA

To test the functional significance of a putative DNA-dependent conformational change, we introduced mutations into the hinge loop, the swivel loop, and at residue D215 to alter the structural flexibility (**Fig. 1C** and **Supplementary** Fig. 2A) and assessed by EMSAs how each substitution affected RexA DNA binding (**Fig. 4**). Point mutations at the glycine hinge (G140A and G140P) have no effect whereas mutating the entire hinge loop to alanines (G140-K143>AAAA) impairs RexA’s ability to bind DNA. Truncation of the swivel loop (Δ239-244, **Supplementary Fig. S2A**) strongly reduces DNA binding, presumably by restraining the movement of the globular domain. In contrast, D215W dramatically enhances DNA binding. D215 lies at a key contact point between the dimerization and globular domains, hydrogen bonding with R261 in the crystallized conformation (**Supplementary Fig. S2A**). Substitution of a bulky tryptophan at this site would introduce a major steric clash (**Supplementary Fig. S2A)**, disrupting the interface between the domains and likely pushing the structure toward an open conformation more readily. None of these changes affect RexA folding or stability (**Supplementary Fig. S6**), suggesting that globular domain movements are necessary to facilitate DNA binding.

**Fig. 4.**
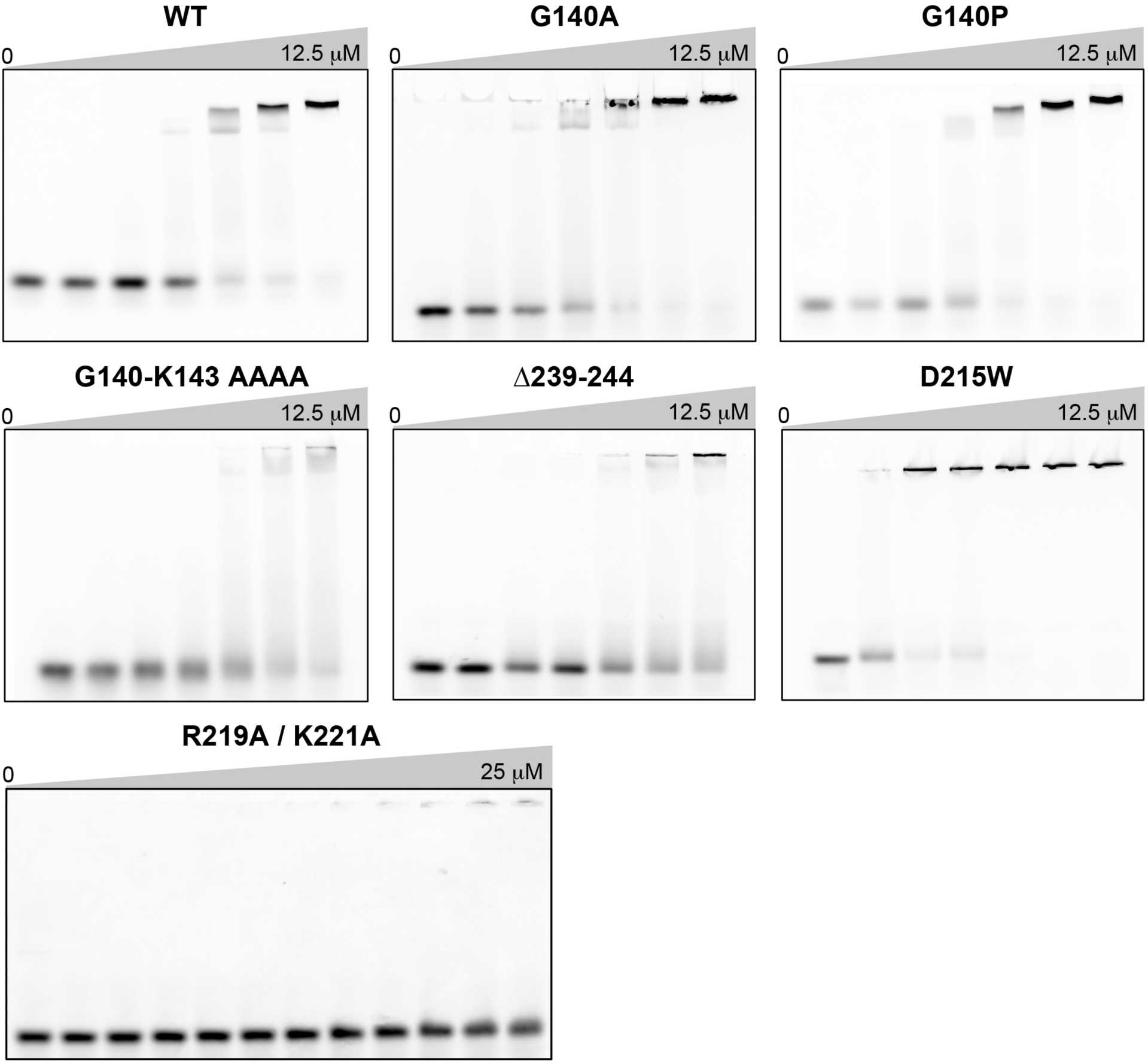
RexA mutants alter DNA binding activity. EMSA analysis of DNA binding by wildtype and mutant RexA proteins. Binding was performed at 25°C for 30 minutes in a 20 μL reaction containing 500 nM of Rex_OR1-OR2 annealed double-stranded DNA with increasing concentrations (0-12.5 μM or 0-25 μM) of each RexA protein construct. Gels were stained with SYBR Gold in 1X TAE buffer for 20 minutes at 25°C to visualize.

Several conserved basic residues line the internal surface of the modeled open conformation of RexA (**Fig. 3B,C and 3J,K**). Among these, residues R219 and K221 at the bottom of the dimerization domain would be oriented into the central cavity, poised to contact DNA (**Supplementary Fig. S2B**). A double mutant substituting alanines at these positions (R219A/K221A) completely abolishes binding to the Rex OR1-OR2 DNA substrate, even at higher concentrations where complete binding is observed for the wildtype protein (**Fig. 4**).

Limited proteolysis is a useful tool to study protein conformational states in that it can define the unstructured and flexible regions of a polypeptide, identify discrete folded fragments that are resistant to cleavage, and monitor changes in protease accessibility as a protein carries out its biological or enzymatic function (72). To further dissect the conformational changes required for RexA DNA binding, we incubated wildtype and mutant RexA constructs with trypsin in the absence or presence of DNA and visualized proteolysis by SDS-PAGE (**Fig. 5A)**. In the absence of DNA, wildtype RexA is almost completely digested by trypsin into small fragments (**Fig. 5A, lane 1 versus control lane C**). Five major undigested fragments appear when DNA is added, suggesting that trypsin’s access to some cleavage sites is limited in this condition (**Fig. 5A, lane 2**). The G140A and G140P mutants, which show no change in DNA binding when analyzed by EMSA (**Fig. 4**), exhibit a pattern of proteolytic cleavage akin to wildtype RexA (**Fig. 5A, lanes 3-6**). The swivel loop truncation (Δ239-344) and R219A/K221A double mutant constructs are proteolyzed to near completion in both conditions (**Fig. 5, lanes 7-8 and 13-14**), reflective of their severe DNA binding defects (**Fig. 4**). Strikingly, we observe stable, undigested fragments with the D215W mutant even when DNA is not present (**Fig. 5, lane 9**). This argues that the proteolytic protection is not simply a consequence of DNA binding and physically blocking access to trypsin sites but rather due to a DNA-induced conformational change that alters protease accessibility and is mimicked by the D215W substitution.

**Fig. 5.**
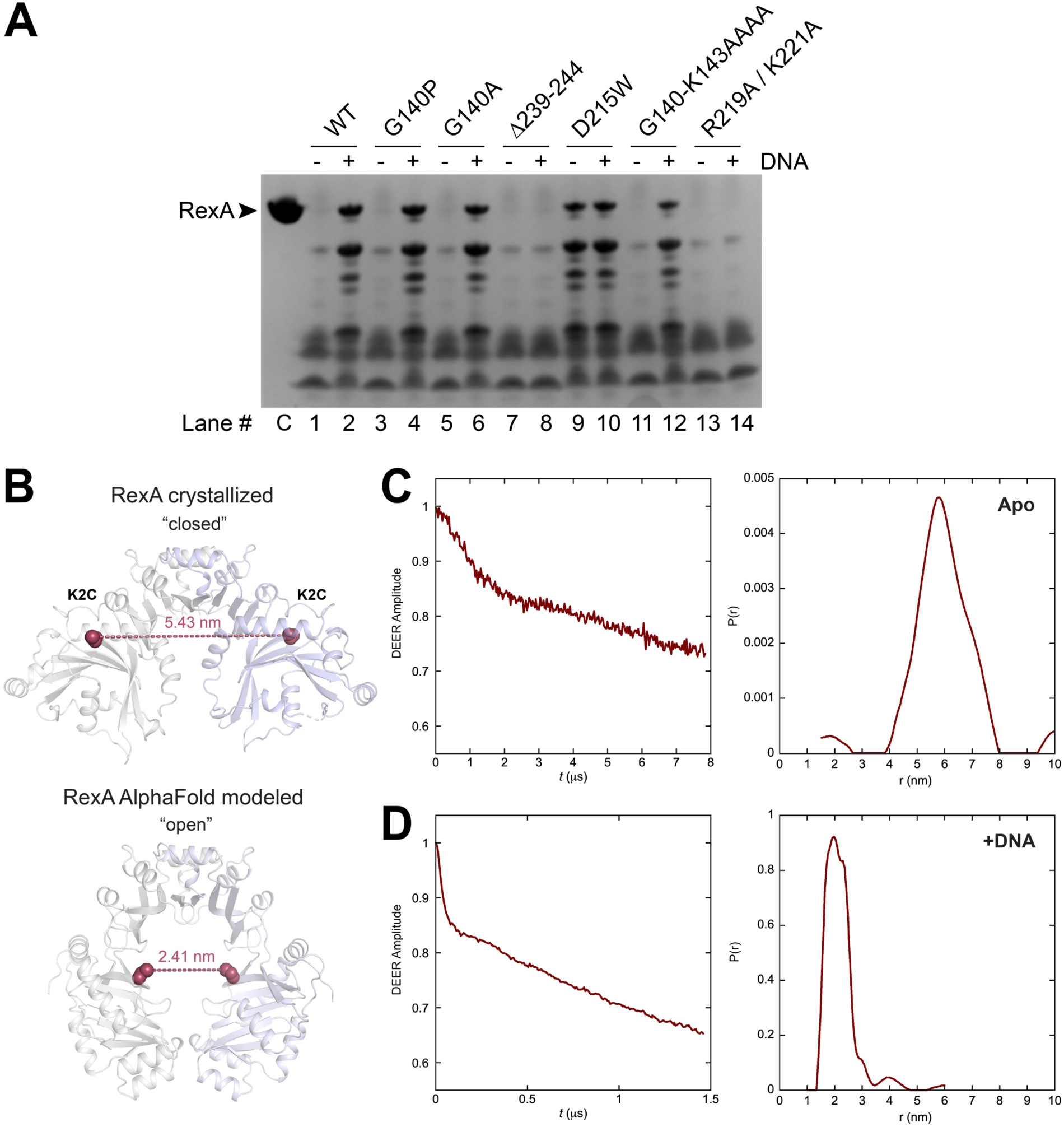
Biochemical and biophysical analyses of RexA mutants support a DNA-dependent conformational change. A. Limited proteolysis of RexA constructs in the absence (-) and presence (+) of 25 μM EMSA_02 (unlabeled) annealed double-stranded DNA. Samples were incubated with 50 μg/mL of trypsin for 30 minutes at room temperature and then subjected to SDS-PAGE and Coomassie staining to visualize. Lanes are numbered below with “C” denoting the untreated control sample (no protease, no DNA). B. Predicted distances between the K2C substitutions (red spheres) in the crystallized (closed) and AlphaFold modeled (open) RexA dimer structures. Individual monomers are colored gray and light blue. C,D. Time domain signals and distance distributions from DEER spectroscopy of K2C RexA in the absence (C) or presence (D) of 20 μM EMSA_02 (unlabeled) annealed double-stranded DNA.

To complement these findings, we further examined DNA-dependent domain movements using <u>P</u>ulse <u>D</u>ipolar ESR <u>S</u>pectroscopy (PDS). PDS techniques are frequently employed to assess different conformational states of proteins and can provide quantitative distances and distance distributions with high concentration sensitivity, yielding critical constraints that can aid in structure refinement and modeling (51, 73–75). The most accessible of these approaches is double electron-electron resonance (DEER) set up as the 4-pulse sequence (76). Distance constraints in the range of ∼1-10 nm can be obtained by measuring the magnitude of the dipolar coupling between the spins of unpaired electrons of nitroxide spin labels and/or metal ions (73, 77–79). Additionally, continuous wave measurements (CW-ESR) can report on short distances from 0.5 to 1.0 nm (80). To facilitate ESR experiments, we mutated the native cysteines in RexA to serines (C258S and C269S) and then introduced a cysteine mutation at lysine 2 (K2C) in the globular domain for labeling with MTSL. Because the RexA dimer is two-fold symmetric, a single cysteine substitution is sufficient to generate a spin label pair. Structure modelling predicts the distance between the K2C cysteine pair to be 5.43 nm in the closed conformation and 2.41 nm in the open conformation (**Fig. 5B**). DEER measurements are consistent with these values, showing a moderate distance distribution centered around 5.8 nm for apo K2C (**Fig. 5C**) and a sharper peak at 2.0 nm in the presence of DNA (**Fig. 5D**).

As a control, we separately introduced a cysteine mutation at residue D168 on the top of the dimerization domain into the C258S/C269S construct. Modelling predicts that the distance between the D186C cysteine pair will be the same in both the closed (apo) and open (DNA bound) conformations (**Supplementary Fig. S7A**). DEER measurements of the MTSL-labeled D168C samples produced traces having only very weak fast-decaying features at early evolution time, indicative of short distances existing below 2 nm in the range where DEER sensitivity is rolling off quickly. This observation was confirmed by double quantum coherence ESR, which showed very fast evolution consistent with strong dipolar couplings and a high likelihood of the distances being at 1 nm or less. Further analysis by X-band CW-ESR revealed that the underlying rigid-limit nitroxide spectrum was broadened ∼2 mT by the large dipolar coupling (**Supplementary** Fig 7B). The broadened part corresponds to close spin label pairs while the rest of the spectra originates from protein molecules containing a spin label on only in one of the protomers, masking the less intense broad spectrum in DEER. Notably, no spectral changes were observed upon DNA binding. The broadened line had an additional wider component exposing the presence of spin-label rotamers and exhibited even larger splitting when the D215W mutation was also incorporated, suggesting that the conformational effects of this mutation can be sensed at the tightly interdigitated region of the dimerization domain. Under the conditions of the ESR experiments, however, D215W does not strictly lock RexA into the open conformation exactly as modeled, which may reflect the intrinsic dynamics of the protein in solution. We thus conclude that while these data support a DNA-dependent conformational change, a more thorough mapping of the λ RexA structure by PDS techniques in the future will be valuable and necessary to delineate the globular domain movements more explicitly and clarify how different mutations can affect different functional states.

### RexA DNA binding does not correlate with T4*rII* exclusion

We next examined how RexA DNA binding and the associated conformational rearrangements we observe *in vitro* contribute to the *in vivo* effects on T4*r*II phage exclusion and modulation of the bistable switch. To study RexA functions *in vivo*, we utilized a previously characterized P_L_P_R_ dual reporter strain (LT1886) wherein the λ immunity region was inserted into the *E. coli lac* operon (21). In this prophage, the P_RM_ maintenance promoter drives expression of the temperature-sensitive *cI857* repressor and the wildtype *rex* genes, the P_R_ lytic promoter drives expression of *cro* and *lacZ*, and the P_L_ lytic promoter drives expression of the firefly luciferase gene *luc* (**Fig. 6A**). Additional reporter strains substituting the RexA R219A/K221A double mutant (LT2294) or the Δ239-244 and D215W conformational mutants (LT2302 and LT2298, respectively) were generated as described in the Material and Methods (58, 59) along with a control strain where the *rexA* and *rexB* genes were absent (*rexAB<>cat*; LT1892) (**Supplementary Table S3**).

**Fig. 6.**
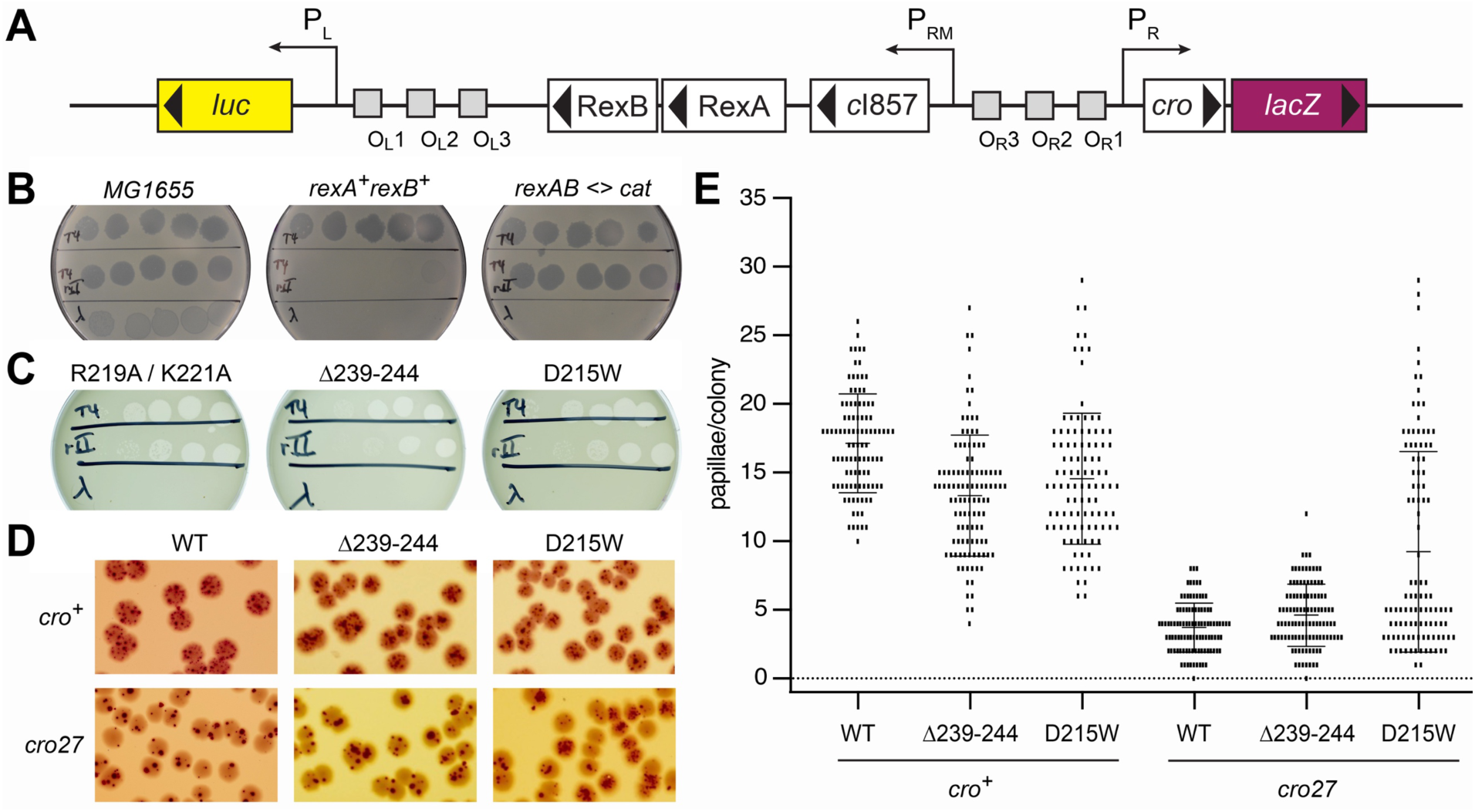
Effects DNA binding and conformational mutants on RexA functions *in vivo*. A. Genetic map of the P_L_P_R_ dual reporter constructed by the insertion of the λ immunity region into the *E. coli lac* operon (21). Reporter strains contain the temperature-sensitive *cI857* repressor with the P_R_ lytic promoter driving expression *lacZ* and the P_L_ lytic promoter driving expression of the firefly luciferase gene *luc*. B,C. Plaque assays testing exclusion of T4, T4rII, and λ*imm* phages (top, middle, and bottom rows on each plate, respectively) by P_L_P_R_ dual reporter strains containing either wildype *rexA* (B) or *rexA* mutants (C). Exclusion phenotypes of *E. coli* strains that either lack the dual reporter insertion (MG1655) or with the *rex* genes replaced with a chloramphenicol resistance cassette (*rexAB <> cat)* are shown for comparison in (B). Strains are labeled as follows (see **Supplementary Table S3** for full details): MG1655, LT351, *rexA^+^rexB^+^*, LT732; *rexAB <> cat*, LT772; R219A / K221A, LT2294; Δ239-244, LT2302; D215W, LT2298. D. Representative papillation from dual reporter strains containing either wildtype *rexA* (WT) or *rexA* mutants Δ239-244 and D215W, respectively, in the context of either *cro^+^* (top row) or *cro27* (bottom) alleles. Strains are as follows (see **Supplementary Table S3** for full details): LT1886, *rexA^+^ cro^+^*; LT1055, *rexA^+^ cro27*; LT2302, *rexA(Δ239-244) cro^+^*; LT2303, *rexA(Δ239-244) cro27*; LT2298, *rexA(D215W) cro^+^*; LT2299, *rexA(D215W) cro27*. E. Quantitation of red papillae in individual colonies for the six genotypes shown in (D). The data are plotted as scatterplots, with each small vertical line indicating the number of papillae found in a single colony. At least 100 hundred colonies were scored for each genotype (WT *cro^+^*, n=100; Δ239-244 *cro^+^*, n=109; D215W *cro^+^*, n=104, WT *cro27*, n=100; Δ239-244 *cro27*, n=106; D215W *cro27*, n=107). The error bars show the SD.

Exclusion of T4*rII* mutant phages is a hallmark of Rex function in λ lysogens (81) and requires both RexA and RexB (28, 82, 83). To study the ability of our reporter strains to exclude this phage, wildtype T4, T4*r*II, and λ phages were spotted onto bacterial lawns and the ability of each phage to grow was determined by the formation of plaques arising from cell lysis and death. All three phages generate plaques on wildtype *E. coli* MG1655 control cells, where the reporter insert carrying the λ immunity region is absent (**Fig. 6B, left**). The *rexA^+^rexB^+^*reporter strain, in contrast, excludes the T4*rII* phage due to Rex function and wildtype λ due to the expression of the CI repressor but permits wildtype T4 phage growth (**Fig 6B, center**), consistent with the behavior of typical λ lysogens. The *rexAB<>cat* strain loses the ability to exclude T4*rII* as the Rex proteins are absent but continues to prevent growth of λ as the *cI857* repressor remains unchanged and able to recognize operator sites in the superinfecting λ phage (**Fig. 6B, right**). Exclusion of T4*rII* is similarly impaired in strains harboring either the RexA R219A/K221A double mutant (**Fig. 6C, left**) or the Δ239-244 swivel loop deletion (**Fig. 6C, center**), both of which disrupt DNA binding *in vitro* (**Fig. 4**). Strikingly, we also observe a T4*rII* exclusion defect in a strain carrying the D215W mutation (**Fig. 6C, right**). Unlike the other RexA mutants analyzed here, D215W enhances DNA binding *in vitro* (**Fig. 4**). This argues against a direct correlation between efficient DNA binding and T4*rII* exclusion activity.

### D251W promotes transition to the lytic state *in vivo*

We previously showed that RexA acts independently of RexB to bias the λ bistable switch toward the lytic state and inhibit lysogeny (24). Using our reporter strains, we investigated whether the RexA conformational mutants Δ239-244 and D215W, which have opposing effects on DNA binding *in vitro*, could also affect lysogenic-to-lytic transitions *in vivo*. Single colonies of the wildtype and each mutant reporter strain were grown on MacConkey Lactose agar at 32-34°C, where they initially appear white but then develop red papillae after several days due to expression of *lacZ* from the P_R_ lytic promoter as cells shift to the lytic state (**Fig. 6D**). The number of red papillae per colony was then scored for at least 100 colonies for each genotype (**Fig. 6E**). For these experiments, reporter strains carrying the non-functional missense mutant allele *cro27* (84) were also constructed and analysed (**Supplementary Table S3, Fig. 6D,6E**) as Rex-dependent effects on colony papillation have been shown to require a functional Cro gene (24). While we observe a comparable distribution of papillation events for each strain in the *cro^+^* background, we find a marked increase in the number and frequency of papillae per colony with the D215W mutant in the context of *cro27* allele compared to wildtype RexA and the swivel loop deletion, both of which show markedly reduced readout from the P_R_ lytic promoter (**Fig. 6E**). This result suggests that the conformational state adopted by D215W, which enhances RexA DNA binding *in vitro*, can also promote transition to the lytic state *in vivo* even when functional Cro repressor is absent and the bistable switch is not locked into the lytic configuration by Cro protein binding to OR3 and repressing P_RM_.

### RexA homologs bind DNA non-specifically

Previous genetic studies identified *rex-*like genes in the temperate Actinobacteriophages Sbash, CarolAnn, and Butters that function as exclusion systems and provide broad defense activity to prevent superinfection by other viruses (85–87). An extensive BLAST search of the Actinobacteriophage Database (36) revealed additional Rex homologs in phages DumpsterDude, Toast, Rubeelu, Blino, and PCoral7 (**Supplementary Figs. S8 and S9**). For a subset of these phages (DumpsterDude, CarolAnn, Toast, Blino, and PCoral7), the genes encoding RexA and RexB homologs are localized in the genome adjacent to a CI-like immunity repressor and Cro-like transcription factor in the same arrangement as they appear in the λ phage immunity region (**Supplementary Figs. S1A and S8**). The orientation of the Rex genes is inverted in other phages (Sbash, Butters, and Rubeelu) with an array of additional genes separating them from the immunity repressor (**Supplementary Fig. S8**). Operator sequences, however, are present between the immunity repressor and the Cro-like transcription factor in these instances, suggesting that the bi-stable switch governing lysogenic-lytic transitions may be similarly regulated in each of these viruses.

To determine if the putative RexA-like proteins present in Actinobacteriophage viruses share the same biochemical properties as λ RexA *in vitro*, we successfully cloned and purified three homologs: Sbash gp30, CarolAnn gp44, and Toast gp42. Each of these, like λ RexA, forms stable dimers in solution when analyzed by SEC-MALS (**Supplementary Fig. S10**). Furthermore, EMSAs reveal that each homolog is also capable of binding DNA substrates containing λ phage OR1-OR2 operator sites, albeit with different affinities (**Fig. 7A**). The overall binding profiles do not change when the sequence of the DNA substrate is scrambled (**Fig. 7B**), mirroring the behavior of λ RexA in filter binding experiments with the same DNA substrates (24). Sbash gp30, CarolAnn gp44, and Toast gp42 also bind DNA substrates that contain their own operator sequences (**Fig. 7C and Supplementary Fig. S11**). Thus, these data indicate that RexA homologs also bind DNA non-specifically.

**Fig. 7.**
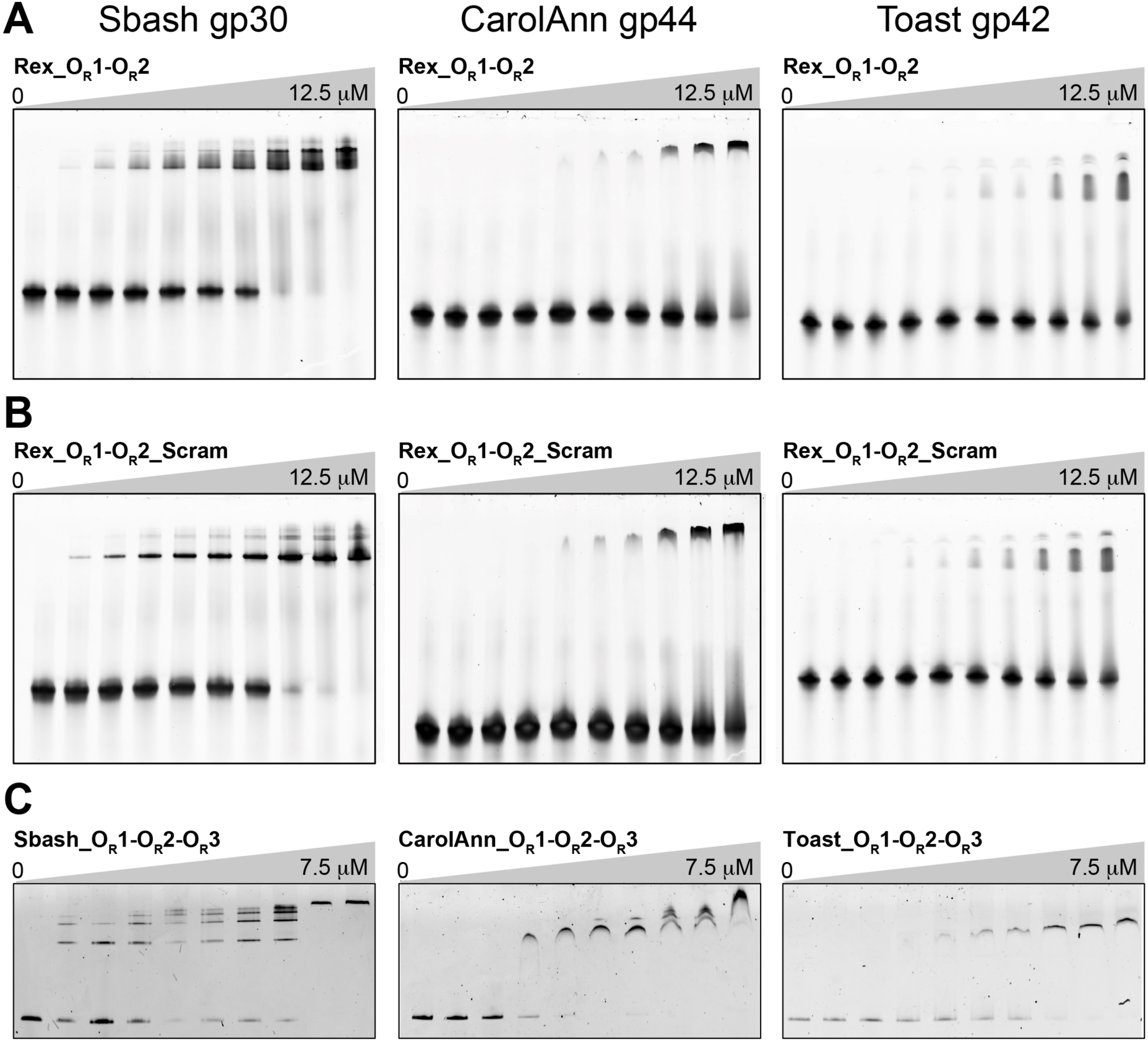
RexA homologs bind DNA non-specifically. EMSA analysis of Sbash gp30, CarolAnn gp44, and Toast gp42 binding to DNA substrates containing λ phage OR1-OR2 operator sites (A), scrambled sequences for λ OR1 and OR2 operators (B), and analogous OR1-OR2-OR3 operator regions specific to each phage (C). Binding was performed at 25°C for 30 minutes in a 20 □L reaction containing 500 nM of each DNA substrate with increasing concentrations (0-12.5 μM or 0-7.5 μM) of each phage protein. Gels were stained with SYBR Gold in 1X TAE buffer for 20 minutes at 25°C to visualize. See **Supplementary Table S2** and **Fig. S11** for detailed descriptions of each substrate.

## DISCUSSION

Here we have shown by X-ray crystallography that the λ RexA protein has a split, two-domain architecture that shares structural homology with RdgC. Both proteins bind DNA non-specifically, suggesting they may have derived from a common ancestor that ultimately diverged into distinct lineages for specialized roles: RexA for phage biology, and RdgC for regulation of recombination. This evolutionary trajectory could explain why some orphan RexA homologs appear in bacterial genomes without an adjacent RexB partner and/or outside the context of an integrated prophage, as is seen in *Shewanella khirikhana* (KEGG ID: STH12_00053) and *Salinisphaera* sp. NP40 (IMG ID: 2790149281).

The domain organization within the RdgC ring-shaped dimer creates a positively charged central pore that is poised to encircle DNA and contact the negatively charged phosphate backbone (67, 69). Despite a shared structural scaffold with RdgC, RexA crystallizes in a closed conformation that is not compatible with this mode of DNA binding (**Figs. 2C and 3A**). Structural superposition and AlphaFold modeling, however, suggest an alternative, open conformation that would permit association with DNA the same manner (**Figs. 2D and 3**). A simple rigid body rotation of the globular domains around G140 in the hinge loop coupled to an unfurling and extension of the swivel loop could facilitate the necessary structural reorganization. Consistent with this, mutations that would constrain conformational flexibility (e.g., G140-K143>AAAA and Δ239-244) reduce RexA DNA binding *in vitro* whereas the steric D215W substitution designed to favor an open conformation enhances DNA binding (**Fig. 4**). We see additional evidence of these predicted changes by limited proteolysis and DEER spectroscopy (**Fig. 5**), which further supports a DNA-dependent conformational change. An analogous rotation of the RdgC center and tip domains has been proposed to enable ring opening and loading of that dimer onto double stranded DNA (69). Domain rotation thus appears to be a conserved mechanistic step needed for DNA binding by both protein families.

We previously established that RexA potentiates prophage induction and can stabilize the lytic configuration of the λ CI-Cro bistable switch, reducing the tendency to return to the immune state (24). Our *in vivo* observations here suggest that RexA’s DNA binding activity and/or conformational state may directly impact this function. Rex-dependent papillation events, which provide a readout of expression from the lytic P_R_ promoter in our reporter system, normally require a functional Cro gene (24) (**Fig. 6D,6E**). The D215W mutant partially overcomes this constraint, producing more colonies with >10 papillation events per colony in the *cro27* background (41/107) compared to wildype RexA (0/100) and the Δ239-244 swivel loop deletion (1/100) (**Fig. 6E**). Wildtype Cro binds to OR3 with high affinity and at high concentrations represses P_RM_ and prevents CI synthesis, which further reinforces the commitment to the lytic state (19, 88, 89). The *cro27* allele, which has a null phenotype (90), contains a missense mutation (G->A) at λ coordinate 38153 that converts an arginine to a glutamine (R38Q) (91). This substitution likely reduces Cro binding to operator DNA as R38 directly contacts the phosphate backbone (92). We speculate that D215W’s enhanced affinity for DNA may allow it to localize to operator sites more efficiently during periods of CI de-repression, perhaps interfering with long-range looping (12), thereby preventing re-establishment of a strongly repressed state.

D215W was engineered to create steric clashing between the globular domains and dimerization domains and generate a more open conformation (**Figs. 2D and 3, Supplementary Fig. S2A**). It is also plausible that the conformational effects associated with this mutation may alter key protein-protein interactions that affect lysogenic-to-lytic transitions. RexB associates with RexA *in vivo* and antagonizes RexA’s modulation of the bi-stable switch (24). Disruption of RexA-RexB interaction would thus be beneficial for lytic transitions while concomitantly preventing phage exclusion (see below). RexA and RexB have both been shown to interact with CI and with Cro by bacterial two-hybrid and RexA has been shown to form stable complexes with larger CI assemblies *in vitro* (24). A more stable interaction with CI dimers might titrate away the repressor and reduce its further oligomerization, which is needed for DNA looping and strong CI repression. Similarly, tighter association with the mutant *cro27* protein product could help stabilize it on DNA and restore disrupted Cro repressor activity.

The Rex system’s association with anti-phage defense traces back to Seymour Benzer’s initial observation that T4 *rII* mutants fail to grow on *E. coli* K12 λ lysogens (81). Wildtype RexA and RexB together exclude T4*rII,* but not wildtype T4 phage (28, 81–83) (**Fig 6B, center**). D215W (enhanced DNA binding), the Δ239-244 swivel loop deletion (impaired DNA binding), and the R219A/K221A double mutant (DNA binding abolished) all lose the ability to exclude T4*rII* despite their radically different DNA binding properties *in vitro* (**Figs. 4 and 6C**). These data argue that RexA’s ability to bind DNA is not a primary determinant of its role in phage exclusion. The failure of these mutants to exclude T4*rII* may instead arise from indirect effects. For example, these mutations may disrupt RexA’s physical association with RexB (23, 24) and/or could alter RexA localization, if DNA interactions are needed to properly position it in the cell for anti-phage defense activities.

Previous genetic studies identified *rex*-like genes in temperate Actinobacteriophages and showed that they, too, can function as exclusion systems, conferring broad immunity against a wide array of other viruses (85–87). Here we demonstrated that purified RexA homologs from the *Mycobacterium* phage Sbash (gp30) and *Gordonia* phages CarolAnn (gp44) and Toast (gp42) all form dimers in solution (**Supplementary** Fig. 10) and can bind double-stranded DNA non-specifically (**Fig. 7**). Although we presently lack atomic-resolution structural data for these proteins, AlphaFold modeling of Toast gp42 predicts with high confidence a two-domain architecture and an assembled dimer arrangement that is reminiscent of our modeled RexA open conformation (**Supplementary Fig. S12**). We note that the genes encoding these RexA-like proteins appear in their respective genomes immediately upstream of a CI-like repressor and a Cro-like transcription factor (**Supplementary Fig. S7**), organized into neighborhoods akin to the λ immunity region (**Supplementary Fig. S1A**). This proximity, coupled with the aforementioned structural and biochemical properties, raises the tantalizing possibility that RexA homologs may also modulate lysogenic-lytic transitions, perhaps independently of their RexB-like partners.

In λ lysogens, RexA’s fine tuning of phage development helps orchestrate entry into the lytic state while its additional role in phage exclusion promotes host fitness. Future studies will establish whether this duality is a generalized feature of other temperate phages or whether it is a specific quirk of λ phage that affords a unique survival advantage.

## DATA AVAILABILITY

The atomic coordinates and structure factors for the bacteriophage λ RexA structure are deposited in the Protein Data Bank with the accession number 8TWQ.

## Supporting information

Adams_RexA_supplementary_info

## ACKNOWLEDGEMENTS

We thank Beth Pennell for continued inspiration and ongoing simulating discussions and the Northeastern Collaborative Access Team (NE-CAT) beamline staff at the Advanced Photon Source (APS) for assistance with remote X-ray data collection.

## AUTHOR CONTRIBUTIONS

M.C.A., C.J.S., L.C.T., D.L.C, and J.S.C. designed the study and analysed data. C.J.H., V.M.J., and H.P. purified and crystallized RexA and collected X-ray diffraction data. C.J.H. and J.S.C. solved the RexA structure and built the model. C.J.S. and J.S.C. carried out computational modelling. M.C.A., C.J.S., and J.S.C. generated and purified all RexA mutants and carried out all biochemical assays. L.C.T. and C.C. performed all genetic experiments. P.P.B. collected and analyzed ESR data with input from J.H.F. M.C.A. and J.S.C. wrote the manuscript with input from L.C.T. and D.L.C.

