## Supplementary material for "The crystal structure of bacteriophage λ RexA provides novel insights into the DNA binding properties of Rex-like phage exclusion proteins": Adams_RexA_supplementary_info

**Supplementary Tables**

Supplementary Table S1. X-ray data collection and refinement statistics.

Supplementary Table S2. Sequences of oligonucleotide substrates.

Supplementary Table S3. *E. coli* K12 strains used for exclusion and papillation assays.

**Supplementary Figures**

Supplementary Fig. S1. Organization and regulation of the λ immunity region. Related to Figs. 1 and 6.

Supplementary Fig. S2. Location of RexA mutants. Related to Figs. 1 and 4.

Supplementary Fig. S3. Structural coordination of bound ions and sulfates. Related to Fig. 1.

Supplementary Fig. S4. Structure and topology of E. coli RdgC. Related to Fig. 2.

Supplementary Fig. S5. Sequence alignment of putative RexA homologs. Related to Figs. 3, 4, and 5.

Supplementary Fig. S6. SEC analysis of RexA mutants. Related to Figs. 4, 5, and 6.

Supplementary Fig. S7. ESR spectroscopy measurements for D168C. Related to Fig. 5.

Supplementary Fig. S8. Gene neighborhoods surrounding RexA-like genes in Actinobacteriophages. Related to Fig. 7.

Supplementary Fig. S9. Sequence alignment of unique RexA-like proteins present in Actinobacteriophage viruses**.** Related to Fig. 7.

Supplementary Fig. S10. SEC analysis of purified RexA homologs. Related to Fig. 7.

Supplementary Fig. S11. Comparison of operator sequences used in binding experiments. Related to Figs. 4, 5, and 7.

Supplementary Fig. S12. AlphaFold model of Toast gp42 and comparison to RexA. Related to Figs. 3 and 7.

**Supplementary Table S1. X-ray data collection and refinement statistics.**

| | RexA SeMet<br>Crystal form 1 | RexA SeMet<br>Crystal form 2<br>PDB: 8TWQ |
|---|---|---|
| **Data collection** | | |
| Space group | P32 2 1 | P32 2 1 |
| Cell dimensions | | |
| *a*, *b*, *c* (Å) | 56.4, 56.4, 326.22 | 55.65, 55.65, 322.12 |
| α, β, γ (°) | 90, 90, 120 | 90, 90, 120 |
| Resolution (Å) | 108.74 – 2.68 (2.81 – 2.68) | 161.06 – 2.05 (2.11 – 2.05) |
| $R_{sym}$ or $R_{merge}$ | 0.127 (0.913) | 0.118 (0.931) |
| $R_{meas}$ | 0.132 (0.997) | 0.128 (1.010) |
| $CC_{1/2}$ (%) | 0.997 (0.634) | 0.998 (0.595) |
| $I$ / σ$I$ | 14.2 (1.7) | 13.4 (1.9) |
| Completeness (%) | 100.0 (100.0) | 99.9 (100.0) |
| Redundancy | 8.9 (8.0) | 6.8 (6.6) |
| **Phasing** | | |
| Initial F.O.M. | 0.3 | |
| Number of sites | 14 | |
| **Refinement** | | |
| Resolution (Å) | | 53.69 – 2.05 |
| No. reflections | | 37768 (3651) |
| $R_{work}$ / $R_{free}$ (%) | | 0.2063 / 0.2512 |
| No. atoms | | 4637 |
| Protein | | 4367 |
| Ligand/ion | | 49 |
| Water | | 221 |
| *B*-factors | | 37.46 |
| Protein | | 37.24 |
| Ligand/ion | | 58.93 |
| Water | | 37.18 |
| R.m.s deviations | | |
| Bond lengths (Å) | | 0.0009 |
| Bond angles (°) | | 1.04 |
| **Ramachandran statistics** | | |
| Favored (%) | | 92.86 |
| Allowed (%) | | 6.04 |
| Outliers (%) | | 1.10 |

*Values in parentheses are for highest-resolution shell. Each dataset was derived from a single crystal.

**Supplementary Table S2. Sequences of oligonucleotide substrates.**

| Oligonucleotide | Sequence |
|---|---|
| EMSA_02_US | 5'- CCACTGGCGGTGAT -3' |
| EMSA_02*_US | 5'- (6-FAM) – CCACTGGCGGTGAT -3' |
| EMSA_02_LS | 5'- ATCACCGCCAGTGG -3' |
| Rex_$O_R$1-$O_R$2_US | 5'- CATTATCACCGCCAGAGGTAAAATAGTCAACACGCACGGTGTTAGAT -3' |
| Rex_$O_R$1-$O_R$2_LS | 5'- ATCTAACACCGTGCGTGTTGACTATTTTACCTCTGGCGGTGATAATG -3' |
| Rex_$O_R$1-$O_R$2_Scram_US | 5'- CATGCCAGGATATTCGCTACAAATAGTACGTGATACGACATCGCGAT -3' |
| Rex_$O_R$1-$O_R$2_Scram_LS | 5'- ATCGCGATGTCGTATCACGTACTATTTGTAGCGAATATCCTGGCATG -3' |
| Toast_$O_R$1-$O_R$2-$O_R$3_US | 5'-<br>CACCGACCCGGCTACCGCTTAACGCGTGTCGTGTCGCACCTGCCGTCTAC<br>CAATGACTACCGGGGATTGTGCCTCATATCCGCAGGCTGCGGAACTACAG<br>GGCTGTAATTTCTTGACGAGCGTCTACCTGCTGCGGAAGAATCATCCGCA<br>GCCTATTGACACCACCCCGTCTACCAGGAGAGACTCATG -3' |
| Toast_$O_R$1-$O_R$2-$O_R$3_LS | 5 -<br>CATGAGTCTCTCCTGGTAGACGGGGTGGTGTCAATAGGCTGCGGATGATT<br>CTTCCGCAGCAGGTAGACGCTCGTCAAGAAATTACAGCCCTGTAGTTCCG<br>CAGCCTGCGGATATGAGGCACAATCCCCGGTAGTCATTGGTAGACGGCA<br>GGTGCGACACGACACGCGTTAAGCGGTAGCCGGGTCGGTG -3' |
| Sbash_$O_R$1-$O_R$2-$O_R$3_US | 5'-<br>CATACGTCGAGTCTGACAGAAACATTCGGCAAACTTTCGAACTGGGTGGC<br>AAACATCATCAGTGTAATTTCCTCCGACTATTGGGTCTTAGGTCAAGAAAC<br>CCCTGGTAGTCCAGGTTTGCCGTTTTAGGAAAAGTTTCGGCATGTCGGTTT<br>GCCGACCCGAATCTACGGTGTATCTTCGGGGATATG -3' |
| Sbash_$O_R$1-$O_R$2-$O_R$3_LS | 5'-<br>CATATCCCCGAAGATACACCGTAGATTCGGGTCGGCAAACCGACATGCCG<br>AAACTTTTCCTAAAACGGCAAACCTGGACTACCAGGGGTTTCTTGACCTAA<br>GACCCAATAGTCGGAGGAAATTACACTGATGATGTTTGCCACCCAGTTCG<br>AAAGTTTGCCGAATGTTTCTGTCAGACTCGACGTATG -3' |
| CarolAnn_$O_R$1-$O_R$2-<br>$O_R$3_US | 5'-<br>CATGGATTGAGGTTCAACCGCCCGTCCTTCGGCACGCAACGCGCAACGG<br>AACCTATGCGGTTACCGCGTTTGCTCGCTCCGGCTTCCTTCTCCGTGTAC<br>CAACTTGTCCCCCGGATTGTGCCGCATACACGCAGGCTGCGGAACTACCG<br>CCGTGTAATTCATTGCCATGCGTGTACCCACCATGCAAGAATCATACGCAG<br>TTGCTTGACACCCAGCCGGTTACCGAGGAGGATCAATGGCGAGCTACACA<br>CGCTGGCCGAAGGGAAGTTGGATG -3' |
| CarolAnn_$O_R$1-$O_R$2-$O_R$3_LS | 5'-<br>CATCCAACTTCCCTTCGGCCAGCGTGTGTAGCTCGCCATTGATCCTCCTC<br>GGTAACCGGCTGGGTGTCAAGCAACTGCGTATGATTCTTGCATGGTGGGT<br>ACACGCATGGCAATGAATTACACGGCGGTAGTTCCGCAGCCTGCGTGTAT<br>GCGGCACAATCCGGGGGACAAGTTGGTACACGGAGAAGGAAGCCGGAGC<br>GAGCAAACGCGGTAACCGCATAGGTTCCGTTGCGCGTTGCGTGCCGAAG<br>GACGGGCGGTTGAACCTCAATCCATG -3' |

**Supplementary Table S3. *E. coli* K12 strains used for exclusion and papillation assays.**

| Strain | Relevant Genotype | Reference/construction |
|---|---|---|
| LT351 | MG1655 | From B. Bochner |
| Cc3LT732 | MG1655 *lacI$^{o}$<>kan-Ter<>luc<>'N pLoL rexB rexA* cI857 *pRoR cro'<>lacZYA*$^{+}$ | Thomason et al., (2019) (1) |
| LT772 | MG1655 *lacI$^{o}$<>kan-Ter<>luc<>'N pLoL (rexB rexA)<>cat* cI857 *pRoR cro'<>lacZYA*$^{+}$ | This work |
| LT1055 | MG1655 *ΔlacI-kan luc-N pLoL rexB*$^{+}$ *rexA*$^{+}$ c*I857ind1 pRoR cro27 cII-lacZYA*$^{+}$ | Thomason et al., (2021) (2) |
| LT1886 | MG1655 *ΔlacI-kan luc-N pLoL rexB*$^{+}$ *rexA*$^{+}$ cI857 *pRoR cro*$^{+}$ *cII-lacZYA*$^{+}$ | Thomason et al. (2021) (2) |
| LT1892 | MG1655 *ΔlacI-kan luc-N pLoL (rexB rexA)<>cat* cI857 *pRoR cro*$^{+}$ *cII-lacZYA*$^{+}$ | Thomason et al. (2021) (2) |
| LT2294 | MG1655 *ΔlacI-kan luc-N pLoL rexB*$^{+}$ *rexA(R219A/K221A)* cI857 *pRoR cro*$^{+}$ *cII-lacZYA*$^{+}$ | This work |
| LT2298 | MG1655 *ΔlacI-kan luc-N pLoL rexB*$^{+}$ *rexA(D215W)* cI857 *pRoR cro*$^{+}$ *cII-lacZYA*$^{+}$ | This work |
| LT2299 | MG1655 *ΔlacI-kan luc-N pLoL rexB*$^{+}$ *rexA(D215W)* cI857 *pRoR cro27 cII-lacZYA*$^{+}$ | This work |
| LT2302 | MG1655 *ΔlacI-kan luc-N pLoL rexB*$^{+}$ *rexA(Δ239-244)* cI857 *pRoR cro*$^{+}$ *cII-lacZYA*$^{+}$ | This work |
| LT2303 | MG1655 *ΔlacI-kan luc-N pLoL rexB*$^{+}$ *rexA(Δ239-244)* cI857 *pRoR cro27 cII-lacZYA*$^{+}$ | This work |

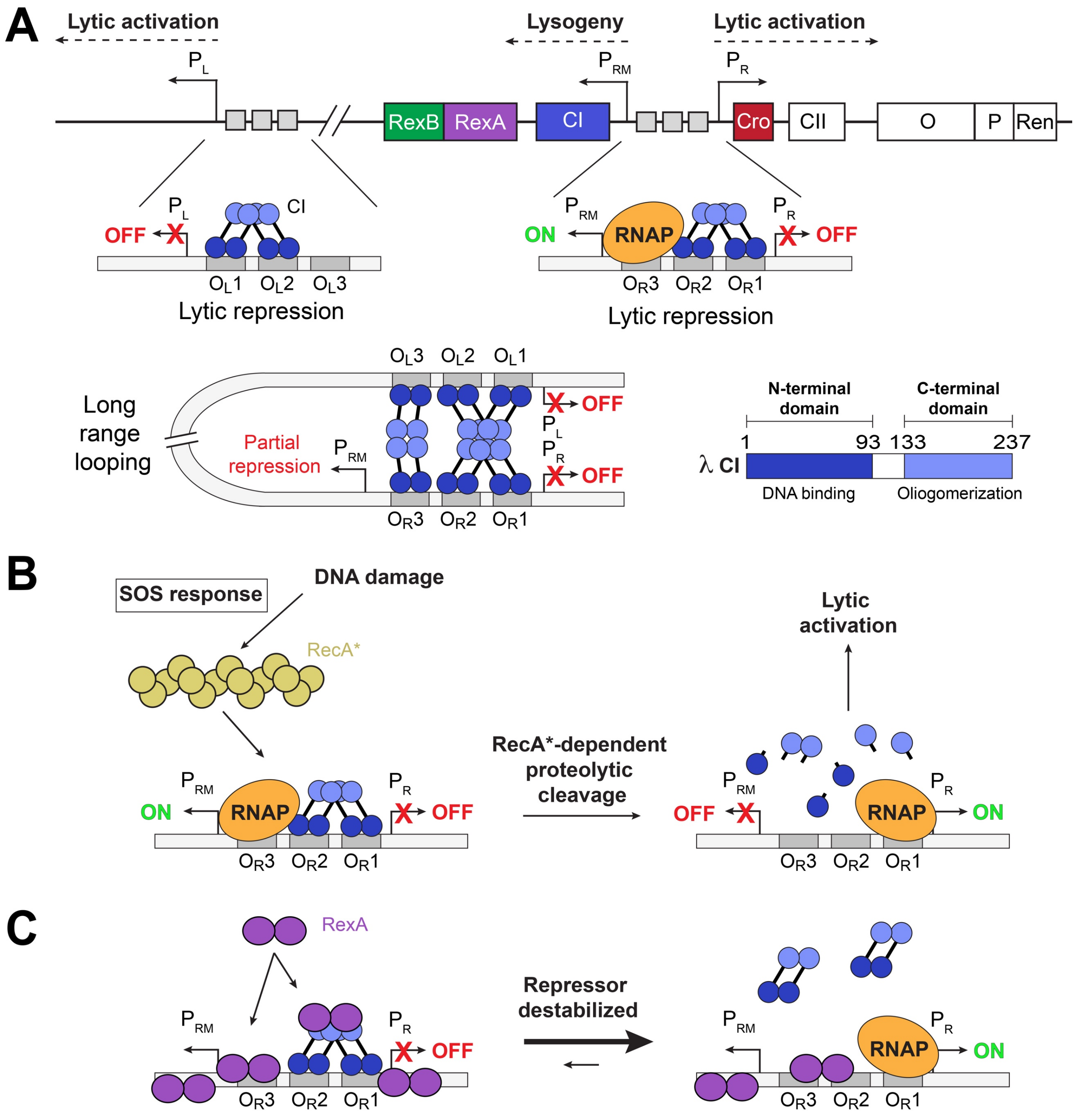


**Supplementary Fig. S1. Organization and regulation of the λ immunity region**. A. Diagram of the λ immunity region. The positions of the $P_L$, $P_{RM}$ and $P_R$ promoters and neighboring genes are labeled. Associated operator sites are marked by gray boxes. In the lysogenic state, CI dimers (blue) are bound cooperatively to OL1 and OL2 and OR1 and OR2 operators to repress the lytic promotors $P_L$ and $P_R$, respectively, and direct transcription from the maintenance promoter $P_{RM}$ via RNA polymerase (RNAP, orange) (3, 4). Long-range DNA looping mediated by the further oligomerization of CI repressor molecules bound to the left and right operators results in stronger repression. Domain organization of CI is shown on the bottom right. B. Prophage induction and lytic activation. Cellular signals like DNA damage and the SOS response activate the RecA protein (RecA*), which promotes proteolytic cleavage of the CI repressor (5). This permits transcription from $P_L$ and $P_R$ lytic promoters. C. Model for RexA modulation of the λ bistable switch. In the absence of RexB, RexA can associate with both the CI repressor (by binding its CTD) and DNA. These protein–protein and protein–DNA interactions may destabilize CI repression and activate the lytic state (2).

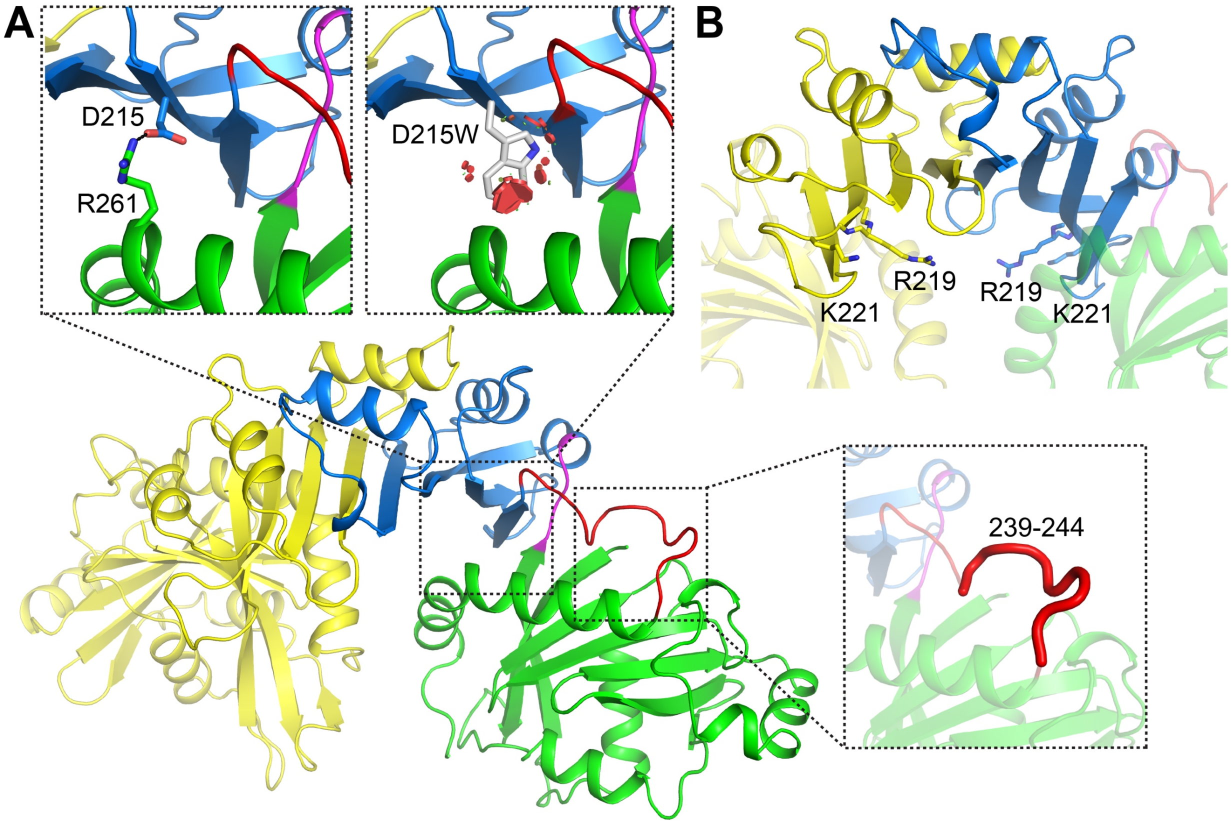


**Supplementary Fig. S2. Location of RexA mutants**. A. Zoomed inserts denote locations of the D215 (blue) and residues 239-244 in the swivel loop (red) that are deleted in the Δ239-244 construct. Modeled D215W mutation (gray) is also shown, with red circles denoting expected steric clashes associated with this substitution. RexA monomers are colored in as in **Fig. 1**. B. Location of R219 and K221 side chains in the dimerization domain.

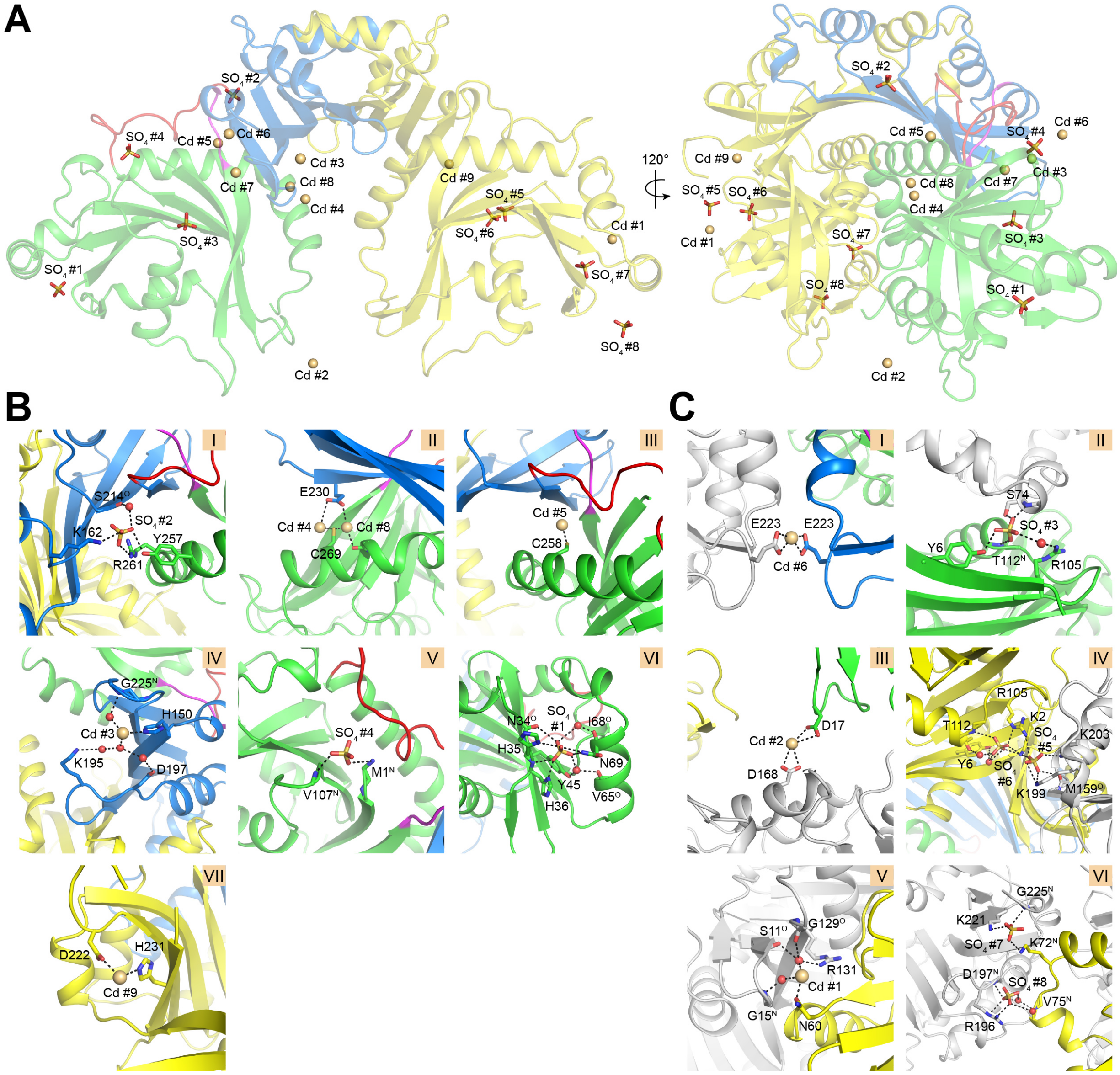


**Supplementary Fig. S3**. **Structural coordination of bound ions and sulfates.** A. Location of ordered sulfates ($SO_4$) and cadmium (Cd) ions associated with the RexA dimer following crystallization. Ions are numbered for reference (see B and C). RexA dimer is colored as in **Fig. 1**. B. Zoomed views intramolecular ionic interactions. Interacting side chains are labeled with hydrogen bonds shown as dashed black lines. Superscripts "N" and "O" denote backbone nitrogen and carbonyl oxygens, respectively. Associated water molecules (red spheres) are shown where applicable. C. Zoomed views of intermolecular ionic interactions mediating crystal contacts. Symmetry-related molecules are colored gray.

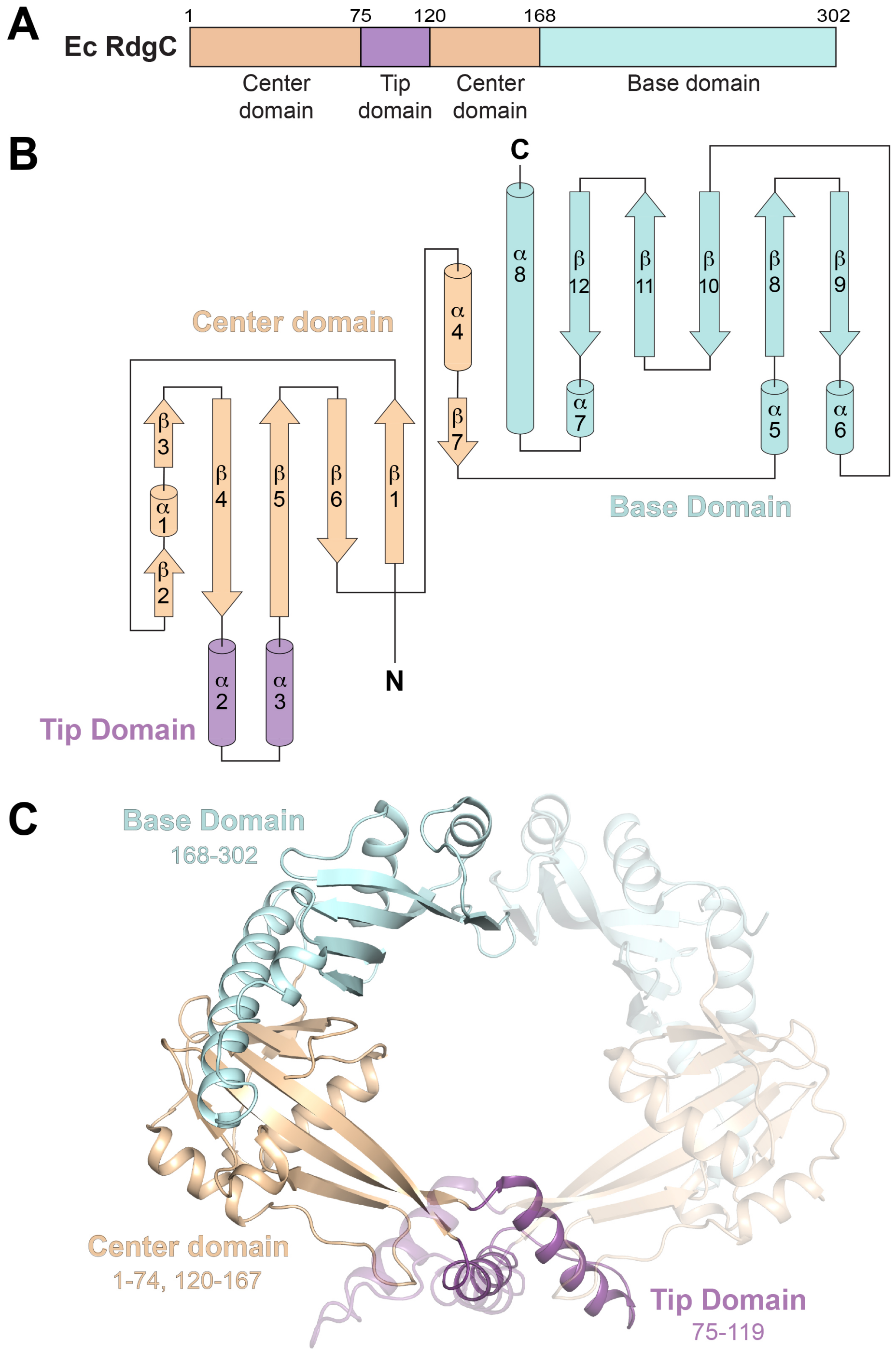


**Supplementary Fig. S4. Structure and topology of *E. coli* RdgC.** A. Domain architecture of *E. coli* (Ec) RdgC. B. Topology diagram of *E. coli* RdgC monomer with coloring as in B. C. Structure of *E. coli* RdgC dimer (PDB: 2OWL). The center, tip, and base domains are colored beige, purple, and cyan respectively in one monomer with accompanying residue numbering. Second monomer is colored the same but rendered partially transparent for contrast.

```
                                                                        β1
BP λ  RexA
BP λ    1  ....................................................MKNGFYATYRSKNK.....  14
BPSfI   1  ....................................................MKNGFYATYRSKNK.....  14
Pbs     1  ....................................................MKNGFYATYRSKNK.....  14
EcK12   1  ....................................................MKNGFYATYRSKNK.....  14
Kp      1  ....................................................MKTGFYAAYRSKDK.....  14
Cd      1  ....................................................MKTGFYAVYQGK.......  12
Kl      1  ....................................................MKTNYFKYDLFPQT.....  14
Ga      1  ....................................................MKISYYKVYEQKENSD.ED  18
Seef    1  ....................................................MKVGFYAAYTETRLN....  15
Er      1  ....................................................MKIGFYAAYAEDG......  13
Yr      1  ....................................................MKMGFYAAYAEAKK.....  14
Mo      1  ....................................................MKLNFHAVYAEDKE.....  14
Ec1     1  ....................................................MKLNFYAVYIQNTS.....  14
Ko      1  ....................................................MKLNFYAVYLQDSK.....  14
Op      1  ....................................................MKVNFYAVYAEDAQ.....  14
Ye      1  ....................................................MKVNFYAVYAEDAK.....  14
Ia      1  ....................................................MKMNFYAAFQQND......  13
Ec2     1  ....................................................MKLNFYAFYQQNG......  13
Seec    1  ....................................................MKMSFYAIYYSNG......  13
Sees    1  MKSPTYGWWKTNRETLIKLQVILFSFRVYHYPYLPLEMCSYDGQSIYITENDMKMSFYAIYYSNG......  65
Cw      1  ....................................................MKMNFYAVYCGMG......  13
Ae      1  ....................................................MKVSFWAYYAKNDI.....  14
Pm2     1  ....................................................MKVSFWAYYAQNTK.....  14
Pm1     1  ....................................................MRVSFWAYYAQDNK.....  14
Mm      1  ....................................................MKVSFLAYYAQDDK.....  14
Pc      1  ....................................................MKINFYAIYSQNQR.....  14
Xb      1  ....................................................MKINFYVFYQENNG.....  14
As      1  ....................................................MKLYYFSYFIRKIS.....  14
Vm      1  ....................................................MKLRYFGYYLREFD.....  14
Fa      1  ....................................................MNVNYFGYYLHDHK.....  14
Wc1     1  ....................................................MRINYYRYYFKYQ......  13
Wc3     1  ....................................................MKINYFKYNFKDGQ.....  14
Ig      1  ......................................MSTEIQNYRKKDLHKTLRYFAYSVADRSYDILS  33
Wc2     1  ..................................................MKKTLRYAAYHFKNTT.....  16
Sk      1  ....................................................MKLTYYSYHFKRN......  13
Ab      1  .............................................MREGVNKLKLSYFTYCFERFD.....  21
Sa      1  ....................................................MKVSYFGYAFERLK.....  14
Ap      1  ....................................................MKVSYFGYSILKQK.....  14
Ha      1  ....................................................MKISYYGYSVERHS.....  14
Gm      1  ....................................................MKVNYFGYCIRNHV.....  14
Pf      1  ................................................MIVSVKISYFGYSLQHKV.....  18
Ps      1  ....................................................MKISYFGYSLQHKV.....  14
                                                               K2
```

```
            β2     α1               β3    TT    β4    TT               β5       α2
BP λ  RexA
BP λ   15  GKDKRSINLSVFLNSLLAD.....NHHLQ..VGSNYLYIHKIDGK..........TFLFTKTNDKSLVQK  67
BPSfI  15  GKDKRSINLSVFLNSLLAD.....NHHLQ..VGSNYLYIHKIDGK..........TFLFTKTNDKSLVQK  67
Pbs    15  GKDKRSINLSVFLNSLLAD.....NHHLQ..VGSNYLYIHKIDGK..........TFLFTKTNDKSLVQK  67
EcK12  15  GKDKRSINLSVFLNSLLAD.....NHHLQ..VGSNYLYIHKIDGK..........TFLFTKTNDKSLVQK  67
Kp     15  GKRKTSLGLLGFIKSLING.....KNSLN..IGNNYLYIHEIDDK..........TFLFTKTNDKSLVQK  67
Cd     13  GKSKLSLGLHDFIHPLLKE.....GNNLK..IGDNYLYIHKIDDK..........TILFTKTNDKSLVQK  65
Kl     15  GQPLATTLQASFL.QLIQNN....TQTVR..LDGAYYSIHHISGD..........VFMFLKTNNTDIIKT  67
Ga     19  KKEKIDINFKDIVDNIIKN.....QHAYS......SEFIAKILKD..........VYVFVKINSTDLIKK  67
Seef   16  KKEKFSINMLKFLMEKSKN.....DNNLN..YGVEYLYVHKINNK..........TFLFTKTKDKNLIKK  68
Er     14  KKNKYSIDVLSFLLQKITQSQ...QISLD..CNDYYLYLHLIDPK..........TILFTKTNDSNLIQK  68
Yr     15  GKLKHEVNLLNFINNKITSNG...GHSLD..CGSHFLYMHRIDDK..........NLLFTKTNDRNLIQK  69
Mo     15  FKKYSIDLKRQINNTIISAK....KYGLE..GSDYNLYMHRISDH..........SFLFTKTNDGELIKK  68
Ec1    15  SLKKQPIDLLSFLTEQVINAK...NHAQQ..IEDYYVFAHHIIND..........SFILTKTFDSELVKK  69
Ko     15  SQSKVSLDLFSAFRTQVIDAK...NHAQQ..IDDYFVFAHHISDA..........TFLLTKTFDSELVKK  69
Op     15  TKKKHPVDLLDIFQNKIIQKK...NHAQA..IEDYYVFAHHIHGE..........CFLVTKTNDSQLVQR  69
Ye     15  TKKKHPVNLMDIFQNKIIKKK...NHAQE..IDDYFVFAHHIHGE..........CFLITKTNDSQLVQR  69
Ia     14  .KEMLPIDLVNGLSELVIKAK...NNALE..VDDYFIFAHHISRE..........CYLITKTYDSDLVKR  67
Ec2    14  .KERFPIDLVSGLSEQVIKAK...NNALE..IDDYFIFAHHISRE..........CYLITKTYDSDLVKR  67
Seec   14  .KDNSPCNLFESLSSQVVEKK...GNALE..VDDYFVFAHHIVDK..........SFLLTKTYDSAFVKR  67
Sees   66  .KDNSPCNLFESLSSQVVEKK...GNALE..VDDYFVFAHHIVDK..........SFLLTKTYDSAFVKR 119
Cw     14  .KVKTPLNLLEQLSQQVIKAK...NNALE..IEDYFIFAHHIVGD..........GFLITKTYDSALVKR  67
Ae     15  TDITSRINLTQAINDKIINKN...SHSLE..SGSDYIYLHPINAQ..........TFLFTKTNDSELIKK  69
Pm2    15  TQGKRSIDLSGIINEKIIDTS...SHAMN..CGDYFTYMHPIDIK..........TFLFTKTSDSDLIKK  69
Pm1    15  TKVRKKIDLSSILDGEIIKKS...NHSLQ..CGDYYIYMHSIDTK..........TILFTKTNDSDLINK  69
Mm     15  TKAKRKVNLDKLFEDLVISKA...SHSFQ..CEDYYTYMHRIDIK..........SFLFTKTNDSDLINK  69
Pc     15  TKDKYKLDLLDLLNKKIIEPK...SHALD..CETHHLYLHRIDDK..........TFLFTKTSDSELIKK  69
Xb     15  IKE..RINLLNLFEENIINPQ...EHKLD..CNTYHLYLHKIDES..........TYLFTKTSDSELIKK  67
As     15  TQEKLIHNISDLLTAYCKSNNNALKGSFK..YGDEKLYLAKIDGY.........KNTYYFLKTNNNDLLKR  74
Vm     15  SQEAFLYNIKPIIDAFVSSNNIELKNSFK..RGDEKLYLTKIAAQ.........KNLYYFVRTSDDDIIKR  74
Fa     15  TNKDYQIDLSDLFDSISNNKYPDINSNLY..YNGDRIYPLPYLGS..........TYLLIQSRDNELIKF  72
Wc1    14  .NKVTFYSIQNLIYQYLHNAPATFQEQFKTTISNDNLYIFETTIN.........NLYFFMRTRDDEVIRK  73
Wc3    15  .NNIIYYNIQNLLNAFTKKAPKKFKSQFT..IFGDNLYLIETTEE.........SLYFFTRTRNDTFIHK  72
Ig     34  NGLRYSINLHAFLECFVEHDDNEFKSSFLD.NNSNNIFLLKTESP.........NVFYFIKTSDGEMIHS  93
Wc2    17  DNENIIFDIRPLLKSYSENGTLKLKSSFE..KSGEKLYLLPAEEEVMDRNRHYTPSLYYFIKTKDQEIIRK  85
Sk     14  .DNKYRHTVLDLLRAYVQNAPAHLKRSFVS.RSGETIFLIPSNNA.........HLFTLITTKNNELIKI  72
Ab     22  DKSQFRTSLRSFIKASTSLEALPFAKQNR..HSGENLYLLPVLEN..........FYLFVQTKDREIIKK  79
Sa     15  DGQKYAFDLRPLLDNFVQWDNKKQKNSFT..HGGEKLFLLKESRG..........FYLFVITRDSDIVKK  72
Ap     15  SDQQYLFDLRPFIRAFVQLDDVKFKSQFS..YADESLFLLPVTDN..........LYLFLMTRTNEIIRK  72
Ha     15  DGEKFLFDLRPFVEAFSKFQNVDFKNRFS..HAGENVYLLRSGKD..........FYLYLITRSQEVIKK  72
Gm     15  TGQKMLFDLRDFLKAYCRHASTAFKNNFF..HNDEHVYLIHHFGD..........TFLFLITRSHELIKK  72
Pf     19  SHETHLVDLSKVLLSFADLDSPTFKSSFR..FNSEYIYLKKLSDN..........VCVLLMTRDGEKFKS  76
Ps     15  SQEAHLVDLSTTILSFADLSSPEFKGSFR..FNSEYVYLKRVSGD..........VCMLLMTRDGERFRS  72
```

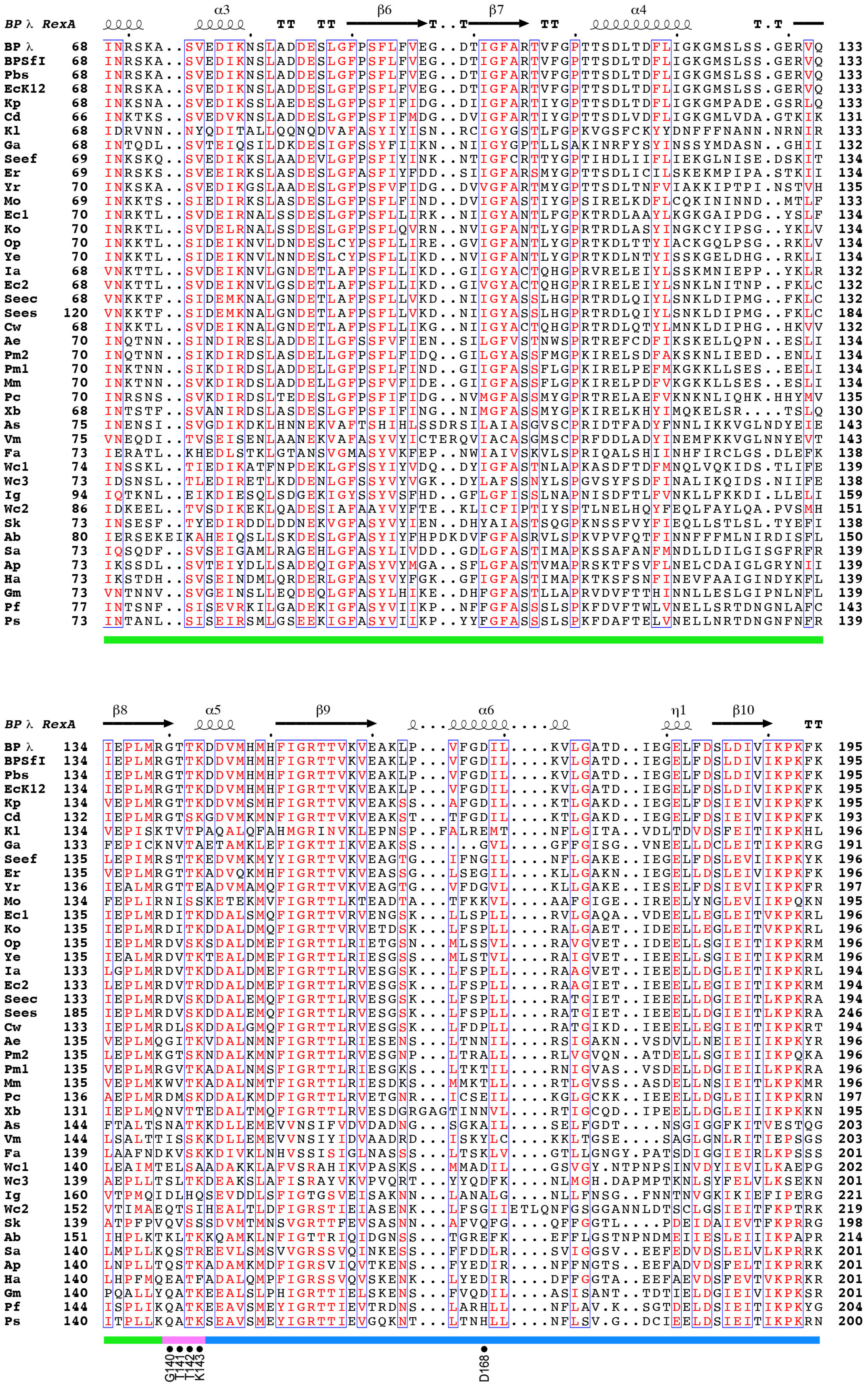

BP λ RexA α3 TT TT β6 T..T β7 TT α4 T.T
BP λ 68 INRSKA..SVEDIKNSLADDESLGFPSFLFVEG..DTIGFARTVFGPTTSDLTDFLIGKGMSLSS.GERVQ 133
BPSfI 68 INRSKA..SVEDIKNSLADDESLGFPSFLFVEG..DTIGFARTVFGPTTSDLTDFLIGKGMSLSS.GERVQ 133
Pbs 68 INRSKA..SVEDIKNSLADDESLGFPSFLFVEG..DTIGFARTVFGPTTSDLTDFLIGKGMSLSS.GERVQ 133
EcK12 68 INRSKA..SVEDIKNSLADDESLGFPSFLFVEG..DTIGFARTVFGPTTSDLTDFLIGKGMSLSS.GERVQ 133
Kp 68 INKSNA..SVEDIKSSLAEDESLGFPSFIFIDG..DIIGFARTIYGPTTSDLTDFLIGKGMPADE.GSRLQ 133
Cd 66 INKTKS..SVEDVKNSLADDESLGFPSFIFMDG..DVIGFARTIYGPTTSDLVDFLIGKGMLVDA.GTKIK 131
Kl 68 IDRVNN..NYQDITALLQQNQDVAFASYIYISN..RCIGYGSTLFGPKVGSFCKYYDNFFFNANN.NRNIR 133
Ga 68 INTQDL..SVTEIQSILDKDESIGFSSYFIIKN..NIIGYGPTLLSAKINRFYSYINSSYMDASN..GHII 132
Seef 69 INKSKQ..SVEDIKSLAADEVLGFPSFIYINK..NTIGFCRTYGPTIHDLIIFLIEKGLNISE.DSKIT 134
Er 69 INRSKS..SVEEIRKSLASDESLGFASFIYFDD..SIIGFARSMYGPTTSDLICILSKEKMPIPA.STKII 134
Yr 70 INKSKA..SVEDIKGSLAADESLGFPSFVFIDG..DVVGFARTMYGPTTSDLTNFVIAKKIPTPI.NSTVH 135
Mo 69 INKKTS..SIEDIRKSLASDESLGFPSFIFND..DVIGFASTIYGPSIRELKDFLCQKININND..MTLF 133
Ec1 70 INRKTL..SVDEIRNALSSDESLGFPSFLLIRK..NIIGYANTLFGPKTRDLAAYLKGKGAIPDG..YSLF 134
Ko 70 INRKTL..SVDELRNALSSDESLGFPSFLQVRN..NVIGFANTLYGPRTRDLASYINGKCYIPSG..YKLV 134
Op 70 INKKTL..SIDEIKNVLDNDESLCYPSFLLIRE..GVIGFANTLYGPRTKDLTTYIACKGQLPSG..RKLV 134
Ye 70 INKKTL..SIDEIKNVLNNDESLCYPSFLLIKD..NVIGFANTLYGPRTKDLNTYISSKGELDHG..RKLI 134
Ia 68 VNKTTL..SVDEIKNVLGNDETLAFPSFLLIKD..GIIGYACTQHGPRVRELEIYLSSKMNIEPP..YKLR 132
Ec2 68 VNKTTL..SVDEIKNVLGSDETLAFPSFLLIKD..GIVGYACTQHGPRIRELEIYLSNKLNITNP..FKLC 132
Seec 68 VNKKTF..SIDEMKNALGNDETLAFPSFLLVKD..NIIGYASSLHGPRTRDLQIYLSNKLDIPMG..FKLC 132
Sees 120 VNKKTF..SIDEMKNALGNDETLAFPSFLLVKD..NIIGYASSLHGPRTRDLQIYLSNKLDIPMG..FKLC 184
Cw 68 INKKTL..SVDEIKNALGNDETLAFPSFLLIKG..NIIGYACTQHGPRTRDLQTYLMNKLDIPHG..HKVV 132
Ae 70 INQTNN..SINDIRESLADDEILGFSSFVFIEN..SILGFVSTNWSPRTREFCDFIKSKELLQPN..ESLI 134
Pm2 70 INQTNN..SIKDIRDSLASDELLGFPSFLFIDQ..GILGYASSFMGPKIRELSDFAKSKNLIEED..ENLI 134
Pm1 70 INKTNN..SIKDIRDSLADDELLGFPSFLFIND..GIIGFASSFLGPKIRELPEFMKGKKLLSES..EELI 134
Mm 70 INKTNN..SVKDIRDSLADDELLGFPSFVFIDE..GIIGFASSFLGPKIRELPDFVKGKKLLSES..ESLI 134
Pc 70 INRSNS..SVKDIRDSLTEDESLGFPSFIFIDG..NVMGFASSMYGPRTRELAEFVKNKNLIQHK.HHYMV 135
Xb 68 INTSTF..SVANIRDSLASDESLGFPSFIFING..NIMGFASSMYGPKIRELKHYIMQKELSR....TSLQ 130
As 75 INENSI..SVGDIKDKLHNNEKVAFTSHIHLSSDRSILAIASGVSCPRIDTFADYFNNLIKKVGLNDYEIE 143
Vm 75 VNEQDI..TVSEISENLAANEKVAFASYVYICTERQVIACASGMSCPRFDDLADYINEMFKKVGLNNYEVT 143
Fa 73 IERATL..KHEDLSTKLGTANSVGMASYVKFEP..NWIAIVSKVLSPRIQALSHIINHFIRCLGS.DLEFK 138
Wc1 74 INSSKL..TIEDIKATFNPDEKLGFSSYIYVDQ..DYIGFASTNLAPKASDFTDFMNQLVQKIDS.TLIFE 139
Wc3 73 IDSNSL..TLEDIRETLKDDENLGFSSYVYVGK..DYLAFSSNYLSPGVSYFSDFINALIKQIDS.NIIFE 138
Ig 94 IQTKNL..EIKDIESQLSDGEKIGYSSYVSFHD..GFLGFISSLNAPNISDFTLFVNKLLFKKDI.LLELI 159
Wc2 86 IDKEEL..TVSDIKEKLQADESIAFAAYVYFTE..KLICFIPTIYSPTLNELHQYFEQLFAYLQA.PVSMH 151
Sk 73 INSESF..TYEDIRDDLDDNEKVGFASYVYIEN..DHYAIASTSQGPKNSSFVYFIEQLLSTLSL.TYEFI 138
Ab 80 IERSEKEIKAHEIQSLLSKDESLGFASYIYFHPDKDVFGFASRVLSPKVPVFQTFINNFFFMLNIRDISFL 150
Sa 73 IQSQDF..SVSEIGAMLRAGEHLGFASYLIVDD..GDLGFASTIMAPKSSAFANFMNDLLDILGISGFRFR 139
Ap 73 IKSSDL..SVTEIYDLLSADEQIGFASYVYMGA..SFLGFASTVMAPRSKTFSVFLNELCDAIGLGRYNII 139
Ha 73 IKSTDH..SVSEINDMLQRDERLGFASYVYFGK..GFIGFASTIMAPKTKSFSNFINEVFAAIGINDYKFL 139
Gm 73 VNTNNV..SVGEINSLLEQDEQLGFASYLHIKE..DHFGFASTLLAPRVDVFTTHINNLLESLGIPNLNFL 139
Pf 77 INTSNF..SISEVRKILGADEKIGFASYVIIKP..NFFGFASSSLSPKFDVFTWLVNELLSRTDNGNLAFC 143
Ps 73 INTANL..SISEIRSMLGSEEKIGFASYVIIKP..YYFGFASSSLSPKFDAFTELVNELLNRTDNGNFNFR 139
BP λ RexA β8 α5 β9 α6 η1 β10 TT
BP λ 134 IEPLMRGTTKDDVMHMHFIGRTTVKVEAKLP...VFGDIL....KVLGATD..IEGELFDSLDIVIKPKFK 195
BPSfI 134 IEPLMRGTTKDDVMHMHFIGRTTVKVEAKLP...VFGDIL....KVLGATD..IEGELFDSLDIVIKPKFK 195
Pbs 134 IEPLMRGTTKDDVMHMHFIGRTTVKVEAKLP...VFGDIL....KVLGATD..IEGELFDSLDIVIKPKFK 195
EcK12 134 IEPLMRGTTKDDVMHMHFIGRTTVKVEAKLP...VFGDIL....KVLGATD..IEGELFDSLDIVIKPKFK 195
Kp 134 VEPLMRGTTKDDVMSMHFIGRTTVKVEAKSS...AFGDIL....KTLGAKD..IEGELFDSIEIVIKPKFK 195
Cd 132 IEPLMRGTSKGDVMKMNFIGRTTVKVEAKST...TFGDIL....KTLGAKD..IEGELFDSIEIVIKPKFK 193
Kl 134 VEPISKTVTPAQALQFAHMGRINVKLEPNSP..FALREMT....NFLGITA..VDLTDVDSFEITIKPKHL 196
Ga 133 FEPICKNVTAETAMKLEFIGKTTIKVEAKSS.......GVL....GFFGISG..VNEELLDCLEITIKPKRG 191
Seef 135 LEPIMRSTTKEDVMKMYYIGRTTVKVEAGTG...IFNGIL....NFLGAKE..IEGELFDSLEVIIKPKYK 196
Er 135 VEPLMRGTTKADVQKMHFIGRTTVKVEASSG...LSEGIL....KLLGAKN..IEGELLDSLEIVIKPKFK 196
Yr 136 IEALMRGTTEADVMAMQFIGRTTVKVEAGTG...VFDGVL....KLLGAKE..IESELFDSIEVIIKPKFR 197
Mo 134 FEPLIRNISSKETEKMVFIGRTTLKTEADTA...TFKKVL....AAFGIGE..IREELYNGLEVIIKPQKN 195
Ec1 135 IEPLMRDITKDDALSMQFIGRTTVRVENGSK...LLSPLL....RVLGAQA..VDEELLEGLEITVKPKRL 196
Ko 135 IEPLMRDITKDDALDMQFIGRTTVRVETDSK...LFSPLL....RALGAET..IDEELLEGLEITVKPKRL 196
Op 135 IEPLMRDVSKSDALDMEFIGRTTLRIETGSN...MLSSVL....RAVGVET..IDEELLSGIEITIKPKRM 196
Ye 135 IEALMRDVTKTEALDMEFIGRTTLRIESGSS...MLSTVL....RAIGVET..IEEELLSGIEITIKPKRM 196
Ia 133 LGPLMRDVTKDDALDMEFIGRTTLRVESGSK...LFSPLL....RAAGVET..IEEELLDGIEITIKPKRL 194
Ec2 133 LEPLMRDVTRDDALDMEFIGRTTLRVESGSK...LFSPLL....RAAGIET..IEEELLEGIEITIKPKRM 194
Seec 133 IEPLMRDVSKDDALEMQFIGRTTLRVESGSK...LFSPLL....RATGIET..IEEELLDGIEITLKPKRA 194
Sees 185 IEPLMRDVSKDDALEMQFIGRTTLRVESGSK...LFSPLL....RATGIET..IEEELLDGIEITLKPKRA 246
Cw 133 IEPLMRDLSKDDALGMQFIGRTTLRVESGSK...LFDPLL....RATGIKD..IEEELLEGIEITIKPKRT 194
Ae 135 VEPLMQGITKVDALNMNFIGRTTLRIESENS...LTNNIL....RSIGAKN..VSDVLLNEIEIIIKPKYR 196
Pm2 135 IEPLMKGTSKNDALKMNFIGRTTLRVESGNP...LTRALL....RLVGVQN..ATDELLSGIEIIIKPQKA 196
Pm1 135 VEPLMRGVTKADALNMSFIGRTTLRIESGKS...LTKTIL....RNIGVAS..VSDELLDGIEIILKPKRA 196
Mm 135 VEPLMKWVTKADALNMTFIGRTTLRIESDKS...MMKTLL....RTLGVSS..ASDELLNSIEITLKPKMR 196
Pc 136 AEPLMRDMSKDDALKMDFIGRTTLRVETGNR...ICSEIL....KGLGCKK..IEEELLDGLEIIIKPRN 197
Xb 131 IEPLMQNVTTEDALTMQFIGRTTLRVESDGRGAGTINNVL....RTIGCQD..IPEELLDGLEIIIKPKKN 195
As 144 FTALTSNATKKDLLEMEVVNSIFVDVDADNG...SGKAIL....SELFGDT....NSGIGGFKITVESTQG 203
Vm 144 LSALTTISSKKDLLEMEVVNSIYIDVAADRD...ISKYLC....KKLTGSE....SAGLGNLRITIEPSGS 203
Fa 139 LAAFNDKVSKKDIVKLNHVSSISIGLNASSS...LTSKLV....GTLLGNGY.PATSDIGGIEIRLKPSSS 201
Wc1 140 LEAIMTELSAADAKKLAFVSRAHIKVPASKS...MMADIL....GSVGY.NTPNPSINVDYIEVILKAEPG 202
Wc3 139 AEPLLTSLTKDEAKSLAFISRAYVKVPVQRT...YYQDFK....NLMGH.DAPMPTKNLSYFELVLKSEKN 201
Ig 160 VTPMQIDLHQSEVDDLSFIGTGSVEISAKNN...LANALG....NLLFNSG..FNNTNVGKIKIEFIPERG 221
Wc2 152 VTIMAEQTSIHEALTLDFIGRSTIEIASENK...LFSGIIETLQNFGSGGANNLDTSCLGSIEITFKPTRK 219
Sk 139 ATPFPVQVSSSDVMTMNSVGRTTFEVSASNN...AFVQFG....QFFGGTL....PDEIDAIEVTFKPRRG 198
Ab 151 IHPLKTKLTKKQAMKLNFIGTTRIQIDGNSS...TGREFK....EFFLGSTNPNDMEIIESLEIIIKPAPR 214
Sa 140 LMPLLKQSTREEVLSMSVVGRSSVQINKESS...FFDDLR....SVIGGSV..EEFEDVDSLELVLKPRRK 201
Ap 140 LNPLLTQSTKADAMKMDFIGRSVIQVTKENS...FYEDIR....NFFNGTS..EEFADVDSFELTIKPRRK 201
Ha 140 LHPFMQEATFADALQMPFIGRSSVQVSKENK...LYEDIR....DFFGGTA..EEFAEVDSFEVTVKPRKR 201
Gm 140 PQALLYQATKEEALSLPHIGRTTIELSKENS...FVQDIL....ASISANT..TDTIELDGIEIVIKPKSR 201
Pf 144 ISPLIKQATKEEAVSMEYIGRTTIEVTRDNS...LARHLL....NFLAV.K..SGTDELDSIEITIKPKYG 204
Ps 140 ITPLLKQATKSEAVSMEYIGRTTIEVGQKNT...LTNHLL....NFLSV.G..DCIEELDSIEITIKPKRN 200
G140 T141 T142 K143
D168

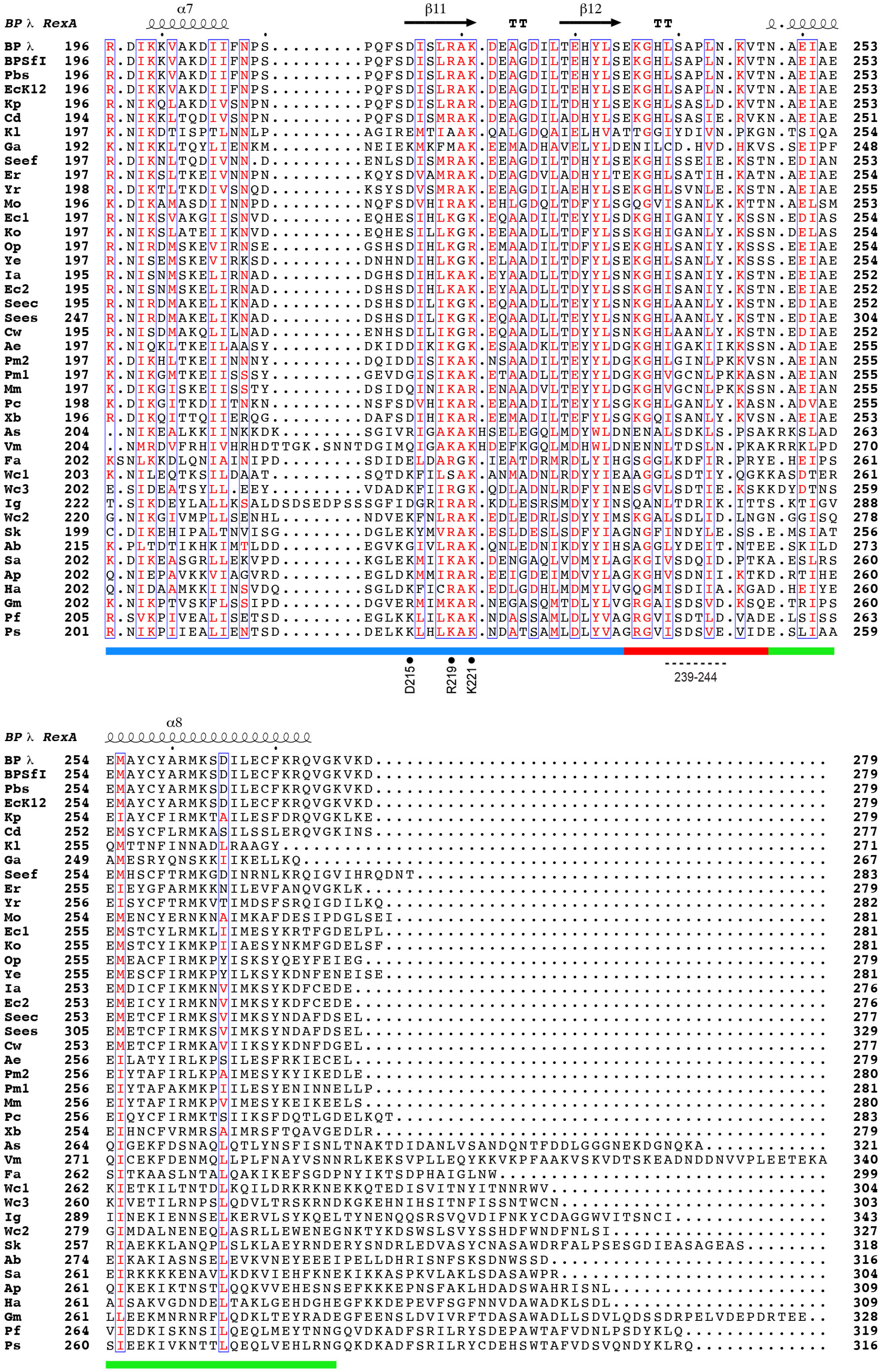

α7 β11 TT β12 TT
BP λ RexA
BP λ 196 R.DIKKVAKDIIFNPS.........PQFSDISLRAK.DEAGDILTEHYLSEKGHLSAPLN.KVTN.AEIAE 253
BPSfI 196 R.DIKKVAKDIIFNPS.........PQFSDISLRAK.DEAGDILTEHYLSEKGHLSAPLN.KVTN.AEIAE 253
Pbs 196 R.DIKKVAKDIIFNPS.........PQFSDISLRAK.DEAGDILTEHYLSEKGHLSAPLN.KVTN.AEIAE 253
EcK12 196 R.DIKKVAKDIIFNPS.........PQFSDISLRAK.DEAGDILTEHYLSEKGHLSAPLN.KVTN.AEIAE 253
Kp 196 R.NIKQLAKDIVSNPN.........PQFSDISLRAR.DEAGDILTDHYLSEKGHLSASLD.KVTN.AEIAE 253
Cd 194 R.NIKKLTQDIVSNPN.........PQFSDISMRAK.DEAGDILADHYLSEKGHLSASIE.RVKN.AEIAE 251
Kl 197 K.NIKDTISPTLNNLP.........AGIREMTIAAK.QALGDQAIELHVATTGGIYDIVN.PKGN.TSIQA 254
Ga 192 K.NIKKLTQYLIENKM.........NEIEKMKFMAK.EEMADHAVELYLDENILCD.HVD.HKVS.SEIPF 248
Seef 197 R.DIKNLTQDIVNN.D.........ENLSDISMRAK.EEAGDILTDHYLSEKGHISSEIE.KSTN.EDIAN 253
Er 197 R.NIKSLTKEIVNNPN.........KQYSDVAMRAK.DEAGDVLADHYLTEKGHLSATIH.KATN.AEIAE 254
Yr 198 R.DIKTLTKDIVSNQD.........KSYSDVSMRAK.EEAGDILAEHYLSEKGHLSVNLE.KSTN.AEIAE 255
Mo 196 K.DIKAMASDIINNPD.........NQFSDVHIRAK.EHLGDQLTDFYLSGQGVISANLK.KTTN.AELSM 253
Ec1 197 R.NIKSVAKGIISNVD.........EQHESIHLKGK.EQAADILTEYYLSDKGHIGANIY.KSSN.EDIAS 254
Ko 197 K.NIKSLATEIIKNVD.........EQHESILLKGK.EEAADLLTDFYLSSKGHIGANLY.KSSN.DELAS 254
Op 197 R.NIRDMSKEVIRNSE.........GSHSDIHLKGR.EMAADILTDFYLSEKGHLSANIY.KSSS.EEIAE 254
Ye 197 R.NISEMSKEVIRKSD.........DNHNDIHLKGK.ELAADILTDFYLSEKGHLSANLY.KSSS.EEIAE 254
Ia 195 R.NISNMSKELIRNAD.........DGHSDIHLKAK.EYAADILTDYYLSNKGHISANIY.KSTN.EEIAE 252
Ec2 195 R.NISNMSKELIRNAD.........DSHSDIHLKAK.EYAADLLTDYYLSSKGHISANIY.KSTN.EEIAE 252
Seec 195 R.NIRDMAKELIKNAD.........DSHSDILIKGK.EQAADLLTEYYLSNKGHLAANLY.KSTN.EDIAE 252
Sees 247 R.NIRDMAKELIKNAD.........DSHSDILIKGK.EQAADLLTEYYLSNKGHLAANLY.KSTN.EDIAE 304
Cw 195 R.NISDMAKQLILNAD.........ENHSDILIKGR.EQAADLLTDYYLSNKGHLAANLY.KSTN.EDIAE 252
Ae 197 K.NIQKLTKEILAASY.........DKIDDIKIKGK.ESAADKLTEYYLDGKGHIGAKIIKKSSN.ADIAE 255
Pm2 197 K.DIKHLTKEIINNNY.........DQIDDIIKIKAK.NSAADILTEYYLDGKGHLGINLPKKVSN.AEIAN 255
Pm1 197 K.NIKGMTKEIISSSY.........GEVDGISIKAK.ETAADLLTEYYLDGKGHVGCNLPKKASN.AEIAN 255
Mm 197 K.DIKGISKEIISSTY.........DSIDQINIKAR.ENAADVLTEYYLDGKGHVGCNLPKKSSN.AEIAN 255
Pc 198 K.DIKGITKDIITNKN.........NSFSDVHIKAR.EEAADILTEYYLSGKGHLGANLY.KASN.ADVAE 255
Xb 196 R.DIKQITTQIIERQG.........DAFSDIHIKAR.EEMADILTEFYLSGKGQISANLY.KVSN.AEIAE 253
As 204 ..NIKEALKKIINKKDK........SGIVRIGAKAKHSELEGQLMDYWLDNENALSDKLS.PSAKRKSLAD 263
Vm 204 ..NMRDVFRHIVHRHDTTGK.SNNTDGIMQIGAKAKHDEFKGQLMDHWLDNENNLTDSLN.PKAKRRKLPD 270
Fa 202 KSNLKKDLQNIAINIPD........SDIDELDARGK.IEATDRMRDLYIHGSGGLKDFIR.PRYE.HEIPS 261
Wc1 203 K.NILEQTKSILDAAT.........SQTDKFILSAK.ANMADNLRDYYIEAAGGLSDTIY.QGKKASDTER 261
Wc3 202 E.SIDEATSYLL.EEY.........VDADKFIIRGK.QDLADNLRDFYINESGVLSDTIE.KSKKDYDTNS 259
Ig 222 T.SIKDEYLALLKSALDSDSEDPSSSGFIDGRIRAR.KDLESRSMDYYINSQANLTDRIK.ITTS.KTIGV 288
Wc2 220 G.NIKGIVMPLLSENHL........NDVEKFNLRAK.EDLEDRLSDYYIMSKGALSDLID.LNGN.GGISQ 278
Sk 199 C.DIKEHIPALTNVISG........DGLEKYMVRAK.ESLDESLSDFYIAGNGFINDYLE.SS.E.MSIAT 256
Ab 215 K.PLTDTIKHKIMTLDD........EGVKGIVLRAK.QNLEDNIKDYYIHSAGGLYDEIT.NTEE.SKILD 273
Sa 202 K.DIKEASGRLLEKVPD........KGLEKMIIKAK.DENGAQLVDMYLAGKGIVSDQID.PTKA.ESLRS 260
Ap 202 Q.NIEPAVKKVIAGVRD........EGLDKMMIRAR.EEIGDEIMDVYLAGKGHVSDNII.KTKD.RTIHE 260
Ha 202 Q.NIDAAMKKIINSVDQ........SGLDKFICRAK.EDLGDHLMDMYLVGQGMISDIIA.KGAD.HEIYE 260
Gm 202 K.NIKPTVSKFLSSIPD........DGVERMIMKAR.NEGASQMTDLYLVGRGAISDSVD.KSQE.TRIPS 260
Pf 205 R.SVKPIVEALISETSD........EGLKKLILKAK.NDASSAMLDLYVAGRGVISDTLD.VADE.LSISS 263
Ps 201 R.NIKPIIEALIENTSD........DELKKLHLKAK.NDATSAMLDLYVAGRGVISDSVE.VIDE.SLIAA 259
D215
R219
K221
239-244
α8
BP λ RexA
BP λ 254 EMAYCYARMKSDILECFKRQVGKVKD................................................ 279
BPSfI 254 EMAYCYARMKSDILECFKRQVGKVKD................................................ 279
Pbs 254 EMAYCYARMKSDILECFKRQVGKVKD................................................ 279
EcK12 254 EMAYCYARMKSDILECFKRQVGKVKD................................................ 279
Kp 254 EIAYCFIRMKTAILESFDRQVGKLKE................................................ 279
Cd 252 EMSYCFLRMKASILSSLERQVGKINS................................................ 277
Kl 255 QMTTNFINNADLRAAGY......................................................... 271
Ga 249 AMESRYQNSKKIIKELLKQ....................................................... 267
Seef 254 EMHSCFTRMKGDINRNLKRQIGVIHRQDNT............................................ 283
Er 255 EIEYGFARMKKNILEVFANQVGKLK................................................. 279
Yr 256 EISYCFTRMKVTIMDSFSRQIGDILKQ............................................... 282
Mo 254 EMENCYERNKNAIMKAFDESIPDGLSEI.............................................. 281
Ec1 255 EMSTCYLRMKLIIMESYKRTFGDELPL............................................... 281
Ko 255 EMSTCYIKMKPIIAESYNKMFGDELSF............................................... 281
Op 255 EMEACFIRMKPYISKSYQEYFEIEG................................................. 279
Ye 255 EMESCFIRMKPYILKSYKDNFENEISE............................................... 281
Ia 253 EMDICFIKMKNVIMKSYKDFCEDE.................................................. 276
Ec2 253 EMEICYIRMKNVIMKSYKDFCEDE.................................................. 276
Seec 253 EMETCFIRMKSVIMKSYNDAFDSEL................................................. 277
Sees 305 EMETCFIRMKSVIMKSYNDAFDSEL................................................. 329
Cw 253 EMETCFIRMKAVIIKSYKDNFDGEL................................................. 277
Ae 256 EILATYIRLKPSILESFRKIECEL.................................................. 279
Pm2 256 EIYTAFIRLKPAIMESYKYIKEDLE................................................. 280
Pm1 256 EIYTAFAKMKPIILESYENINNELLP................................................ 281
Mm 256 EIYTAFIRMKPVIMESYKEIKEELS................................................. 280
Pc 256 EIQYCFIRMKTSIIKSFDQTLGDELKQT.............................................. 283
Xb 254 EIHNCFVRMRSAIMRSFTQAVGEDLR................................................ 279
As 264 QIGEKFDSNAQLQTLYNSFISNLTNAKTDIDANLVSANDQNTFDDLGGGNEKDGNQKA........... 321
Vm 271 QICEKFDENMQLLPLFNAYVSNNRLKEKSVPLLEQYKKVKPFAAKVSKVDTSKEADNDDNVVPLEETEKA 340
Fa 262 SITKAASLNTALQAKIKEFSGDPNYIKTSDPHAIGLNW.................................... 299
Wc1 262 KIETKILTNTDLKQILDRKRKNEKKQTEDISVITNYITNNRWV............................... 304
Wc3 260 KIVETILRNPSLQDVLTRSKRNDKGKEHNIHSITNFISSNTWCN.............................. 303
Ig 289 IINEKIENNSELKERVLSYKQELTYNENQQSRSVQVDIFNKYCDAGGWVITSNCI................... 343
Wc2 279 GIMDALNENEQLASRLLEWENEGNKTYKDSWSLSVYSSHDFWNDFNLSI......................... 327
Sk 257 RIAEKKLANQPLSLKLAEYRNDERYSNDRLEDVASYCNASAWDRFALPSESGDIEASAGEAS........ 318
Ab 274 EIKAKIASNSELEVKVNEYEEEIPELLDHRISNFSKSDNWSSD............................... 316
Sa 261 EIRKKKKENAVLKDKVIEHFKNEKIKKASPKVLAKLSDASAWPR.............................. 304
Ap 261 QIKEKIKTNSTLQQKVVEHESNSEFKKKEPNSFAKLHDADSWAHRISNL......................... 309
Ha 261 AISAKVGDNDELTAKLGEHDGHEGFKKDEPEVFSGFNNVDAWADKLSDL......................... 309
Gm 261 LLEEKMNRNRFLQDKLTEYRADEGFEENSLDVIVRFTDASAWADLLSDVLQDSSDRPELVDEPDRTEE.. 328
Pf 264 VIEDKISKNSILQEQLMEYTNNGQVDKADFSRILRYSDEPAWTAFVDNLPSDYKLQ.............. 319
Ps 260 SIEEKIVKNTTLQEQLVEHLRNGQKDKADFSRILRYCDEHSWTAFVDSVQNDYKLRQ............. 316

**Supplementary Fig. S5. Sequence alignment of putative RexA homologs**. Sequence alignment of RexA homologs with the secondary structure of the bacteriophage  RexA mapped above. Colored bars beneath the alignment denote structural segments as follows: globular domain, green; dimerization domain, blue; hinge loop, magenta; swivel loop, red (see **Fig. 1**). Positions of conformational mutations are marked below (see **Figs. 4 and 5** and **Supplementary Fig. S3**). Sequence shading indicates conservation: white text on red background, 100% conserved; boxed red text on white background, 70% conserved. Abbreviations are as follows with accompanying NCBI accession numbers or KEGG IDs (6) : BPλ, bacteriophage λ RexA (vg:3827058); BPSfI, bacteriophage SfI gp47 (vg:24722216); Pbs, *Paenibacillus sonchi* (pson:JI735_34930), EcK12, *Escherichia coli* K-12 BW2952 (ebw:BWG_3702), Kp, *Klebsiella pneumoniae* (WP_117261707.1); Cd, *Cedecea davisae* (WP_202303730.1); Kl, *Klebsiella* multispecies (WP_071995728.1); Ga, *Gilliamella apicola* (WP_065635067.1); Seef, *Salmonella enterica* subsp. enterica FNW19H96 (EDW1732907.1); Er, *Erwinia* sp. S38 (WP_200545456.1); Yr, *Yersinia ruckeri* (WP_234057212.1); Mo, *Morganellaceae* multispecies (WP_154640079.1); Ec1, Escherichia coli (WP_253764069.1); Ko, unclassified *Kosakonia* multispecies (WP_200133690.1); Op, *Obesumbacterium proteus* (WP_234559868.1); Ye, *Yersinia enterocolitica* (WP_050128085.1); Ia, *Izhakiella australiensis* (WP_096777782.1); Ec2, *Escherichia coli* (WP_197940034.1); Seec, *Salmonella enterica* subsp. *enterica* serovar Cubana (seec:CFSAN002050_11670); Sees, *Salmonella enterica* subsp. *enterica* serovar Saintpaul (ECA2934630.1); Cw, *Citrobacter werkmanii* (WP_085048607.1); Ae, Arsenophonus endosymbiont of Apis mellifera (aet:LDL57_11880); Pm2, *Proteus mirabilis* (WP_143474652.1); Pm1, *Proteus mirabilis* (WP_206081156.1); Mm, *Morganella morganii* (WP_049246396.1); Pc, *Photorhabdus cinerea* (WP_166310405.1); Xb, *Xenorhabdus bovienii* (WP_038244014.1); As, *Alteromonas stellipolaris* R10SW13 (aaw:AVL56_04330); Vm, Vibrio mimicus (vmi:AL543_00300); Fa, *Frateuria aurantia* (fau:Fraau_1963); Wc1, *Wohlfahrtiimonas chitiniclastica* (WP_213398763.1); Wc3, *Wohlfahrtiimonas chitiniclastica* (WP_094493134.1); Ig, Ignatzschineria sp. HR5S32 (ign:MMG00_12050); Wc2, *Wohlfahrtiimonas chitiniclastica* (WP_213405574.1); Sk, *Shewanella khirikhana* (skh:STH12_00053); Ab, *Acinetobacter baumannii* BJAB0715 (abab: BJAB0715_02483); Sa, *Salinisphaera* sp. (MBS61511.1); Ap, *Abyssibacter profundi* (WP_109719971.1); Ha, *Halomonas* sp. 3F2F (WP_226930571.1); Gm, *Gallaecimonas mangrovi* (WP_115720386.1); Pf, *Pseudomonas fragi* (pfz:AV641_12615); Ps, *Pseudomonas syringae* (WP_198722127.1).

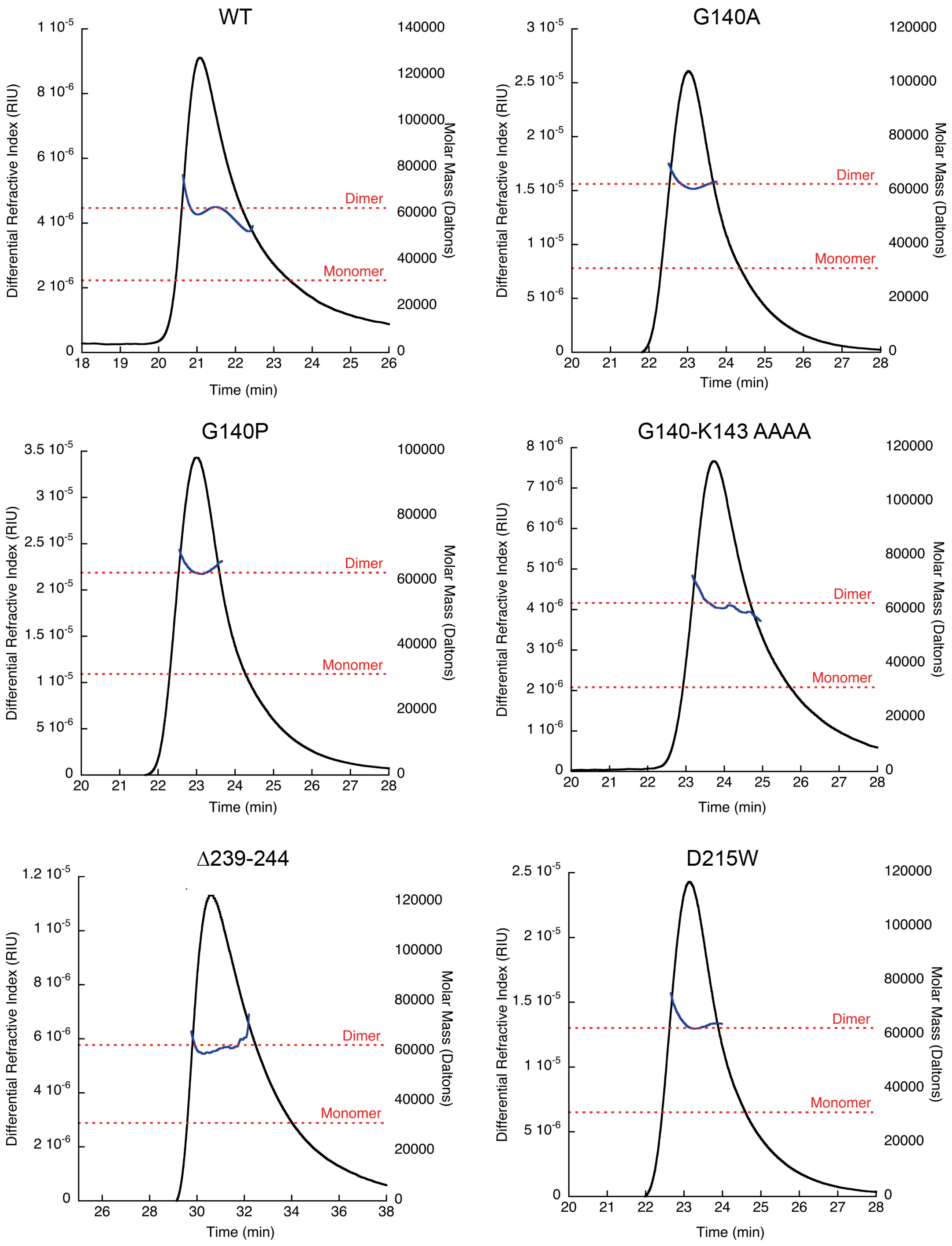


**Supplementary Fig. S6. SEC analysis of RexA mutants**. Black line denotes differential refractive index and blue line denotes measured mass across each peak. Dashed red lines indicate the predicted molecular weight of a RexA monomer and dimer.

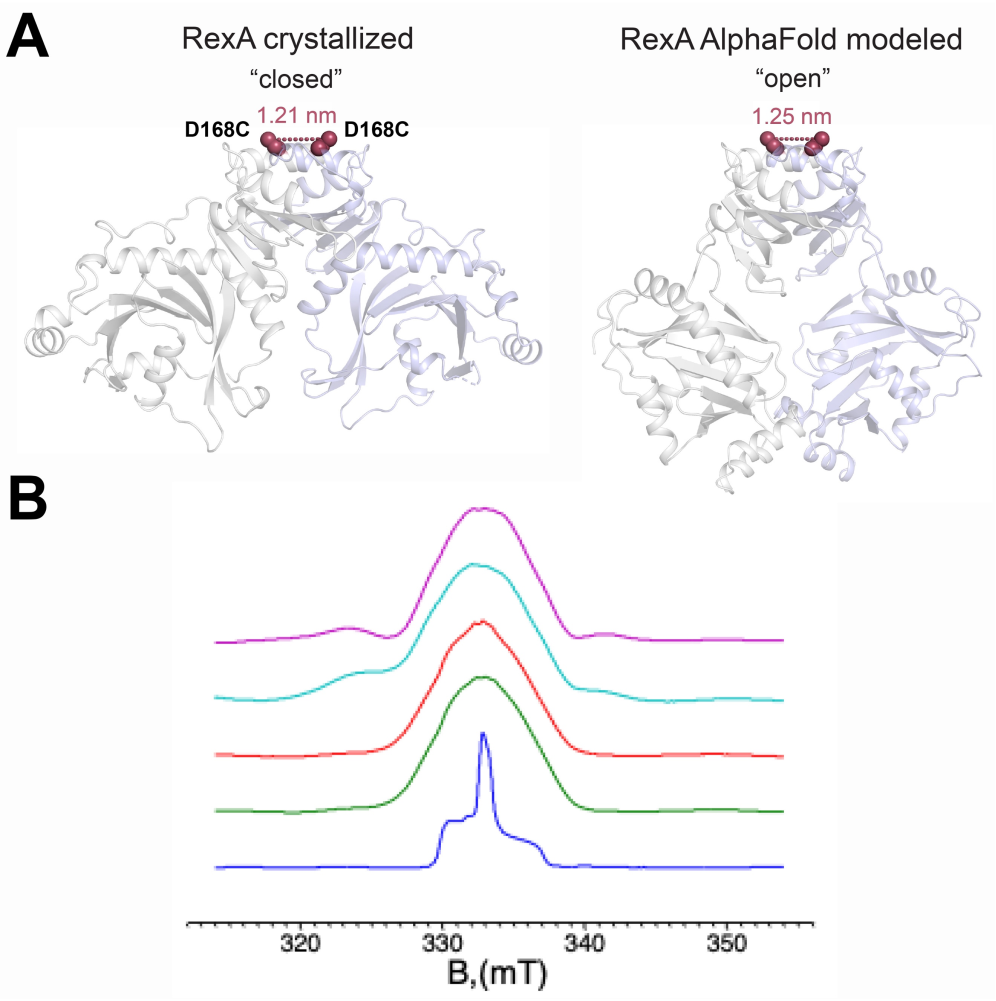


**Supplementary Fig. 7. ESR spectroscopy measurements for D168C.** A. Predicted distances between the D168C substitutions (red spheres) in the crystallized (closed) and AlphaFold modeled (open) RexA dimer structures. B, Integrated CW ESR spectra of D168C (green); D168C/DNA (red); D128C/D215W (cyan); and D168C/D215W/DNA (magenta) spin-labeled RexA mutants. The rigid-limit unbroadened nitroxide spectrum (blue) was plotted as a reference. All D168C spectra show large dipolar broadening ~2 mT corresponding to ~1 nm distance between spin-labels. Spectral features caused by nitroxide labels not having partners were subtracted out. There is no visible effect of DNA but D215W mutation adds shoulders to the spectra.

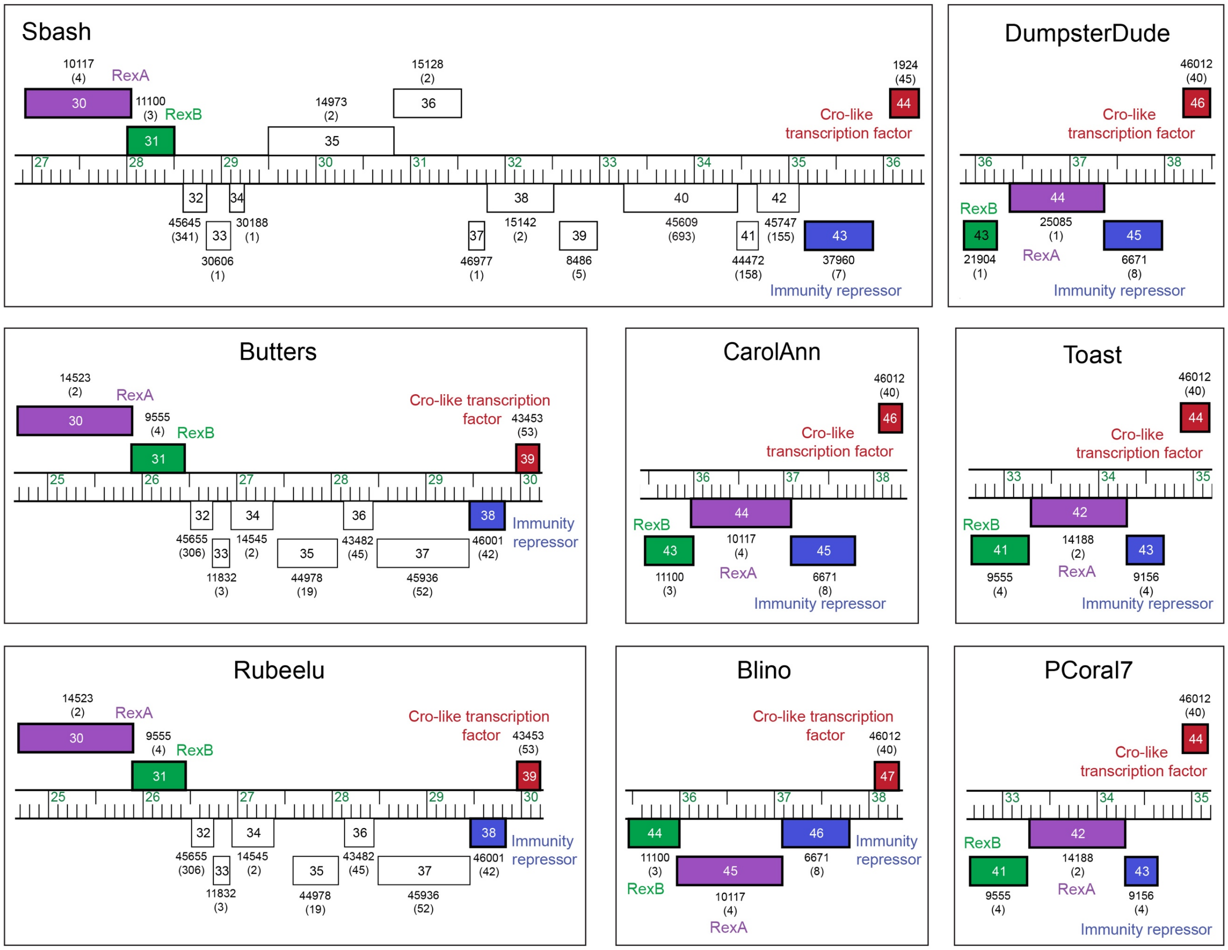


**Supplementary Fig. S8. Gene neighborhoods surrounding RexA-like genes in Actinobacteriophages.** Genomic organization around RexA-like genes in Sbash, DumpsterDude, Butters, Rebeelu, CarolAnn, Blino, Toast, and PCoral7 phages generated via Phamerator (7). Individual genes are depicted as boxes with the gene name inside the box and the Phamily designation and number of Pham members (in parentheses) shown below each box. Genes corresponding to RexA (purple), RexB (green), a CI-like immunity repressor (blue), and Cro-like transcription factor (red) are colored and labeled.

```
λ RexA              1 : --------MKNGFYATYRSKNKGKDKRSINLSVFLNSLLADNH-HLQVGSNYLYIHKIDG-------KTFLFTKTNDKSLVQKINRSK :  72
Sbash gp30          1 : -------------------MTGERQIVAYGLRVEVDGRAQRRTKNPPPHTPYPANFNGAGADLLMFFSSFVNALPTD----------- :  58
CarolAnn gp44       1 : -----------------MGTWGDRRIVLYTLRVALDGRVFRTS--AAAGEPFVANFNGSGADLLHFLSSFIEALPTD----------- :  58
Toast gp42          1 : -------------------------MPSYGIRFYKVELFNGNK-----QTPLSFIE-EDGGKSWRYGE-HLATVLE------------ :  44
DumpsterDude gp44   1 : --------------------MPRRTMTVTEVWAHPVG--SRKE-----KDRVDLTALPDGMDLLHAFYGFAADVDAR----------- :  50
Butters gp30        1 : MLWDRTSHVLSISYANLRTSTHPPLMTNYGFRLFQVGVYRNGR-----GDPCAVDA--IGDDKQHFRD-YLFDLAN------------ :  68

λ RexA             73 : ASVEDIKNSLADDESLGFPSFLFVEGDTIGFARTV-----------------------------FGPTTSDLTDFLIGKGMSLSSGER : 131
Sbash gp30         59 : ------KLIRRGDRHFGIPEEVERLGRTI-------------------------RLRLSGGESGRRSKIT------------------ :  97
CarolAnn gp44      59 : ------KHIERDERHFGQPTSIRRRGRTI-------------------------SWRMDGGESGRKSQIR------------------ :  97
Toast gp42         45 : --------RFKENQVRGFPRSQTPDGESVDQ--------------------IRKKTAMVFNAVDRRGPNTVVAEYRVGRSDDFDR--- : 101
DumpsterDude gp44  51 : ------QLLKRENESFASVVSLEEKGRAV-------------------------TLAVDVGRFGERGTIT------------------ :  89
Butters gp30       69 : --------RKKDCVVHGSPPRSKEEGDSDESADVDDHESVDDDAEQQPTGAKRPRPVLVITDVSFRGD-HVVIEYLYGRDLGYTH--- : 144

λ RexA            132 : VQIEPLMRGTTKDDVMHMHFIGRTTVKVEAKLPVFGDILKVLGATDIEGELFDSL---DIVIKPKFKRDIKKVAKDIIF--------- : 207
Sbash gp30         98 : ------LKAGGKEETREPSGVEWEPFRVAIVIPKDFNQ----GWLLVEKSGYHSV---PTEWRQELVRAFKAAYPDYVL-KISSITEM : 171
CarolAnn gp44      98 : ------LRRDADAQERDRSGVEWEPFWAFAVVPENSVV----GWLLVEKAGRHSL---PTEWRKELQRQFAGAYRGYRL-EIGIVREM : 171
Toast gp42        102 : --AYPAPDLETSDYLELNGYAPARPYRAVLMVPPEGEV----GMLAVEAIGRTCPYEFFTKWATKWSRDYQAELD---SSDPLDDDKV : 180
DumpsterDude gp44  90 : -----DVTTMIERDRFTRTDAISVITHGIFTLPKGCTS----ALVFIERSGNQSG---IIRVLELFQHQFRLAYPDLIL-ETTAVVES : 164
Butters gp30      145 : --AGYSEQDTTAKAVRIANLKSVRPYRSVFMFPDVGNS----GVVAVEDASRAHASKKLEQWLKAWAREEAHAVAEIKRKETGKKSIK : 226

λ RexA            208 : ------NPSPQFSDISLRAKDEA----GDILTEHYLSEKGHLS--APLN---------KVTNAEIAEEM--AYCYARMK--------- : 263
Sbash gp30        172 : SLWSQVE--NSN-EPRLLAVEVIMRSS----DSSGDHSAGHPANMATYDTHVWEAVHAQPG--RVLRQLRRRFTRRKTK---DGL--- : 244
CarolAnn gp44     172 : SLWAQVE--NALDEERLIGFEVAYRSA---STTAGSGAAGYERGMEKYERRMWRAPADQPKKGRKLREFRRAHT-KTLI---EGT--- : 247
Toast gp42        181 : RAWWRLKPTPLGSKSQLVDFIKGGTIEELILINHYIDKTRNER-QERFR------------VKAKVRTGQRPWAKKRLNEV----FEV : 251
DumpsterDude gp44 165 : EAWLSYA--------SLVKVSAHRRA-------SGSDEA--DNNSKRIKRSEYGELAHTLVPARGERTLPR-FLYDRLV---AGA--- : 228
Butters gp30      227 : PVWWSMRFTPLSDPERLAALMKNGTSSKIVLTKQGGSDARTPG-RSPLK---VEMGLDEPASIAKARRLITGWLPKFNRTPQAGDAQV : 310

λ RexA            264 : -----------------------SDILECFKRQVGKVKD------------------------------------------------- : 279
Sbash gp30        245 : ---------VEITQPLDVDDLSDDDAKIELKDDVRQIKARVLNSEGRKKTVIFGGRE-PTMTIVMDGVFDLSPSATKFWQEARSA--- : 319
CarolAnn gp44     248 : ---------REIELPFEADDLSDDRYTIRLRDDVAEIKATLYNSEGQPKTVVFEGLD-PQQTIVMEDTALAPPDQDRFDTECRSA--- : 322
Toast gp42        252 : DSDEVFARQLAENFGNSVEHLDIDDGYVVVRTEA-GVKNIS--PSRMPEVFTYPVSD----------DR--PT-MEEFLTAV------ : 317
DumpsterDude gp44 229 : ---------LKPSELLDFDE-SDE---------AAHVE-VTLEHNGQRKTFLLGHEKRPSISYLL------SNHGEDEWSLERVRDYA : 290
Butters gp30      311 : AANRELAAVLADNYQ-GFDEEDYDDAWIEVKDAAGKAKKIS--PSRWADIFIYPVRN----------STECPP-PALFYRRV------ : 378

λ RexA              - : --------------------------------------------------------- :   -
Sbash gp30        320 : ----VVDLATSGGVSLAPKWDTGEWEHPENAIKVEVSLTDEPDQDDAASAGGTGTTA : 372
CarolAnn gp44     323 : ----VRDLAASNMVGLAPKWDTGDWVHPEDAVKLEVKSDAPSPERDQGSEAN----- : 370
Toast gp42        318 : -KRHAIPLAKVID----AQVDFDSW-------------------------------- : 337
DumpsterDude gp44 291 : FDTHADTIYGRLGWEWTPAHTVGAWTDSQRGARLVVRGGEQS--------------- : 332
Butters gp30      379 : -QESVRPLEKSLE----LSIDWAGWGGDSG--------------------------- : 403
```

**Supplementary Fig. S9. Sequence alignment of unique RexA-like proteins present in Actinobacteriophage viruses.** Alignment includes unique RexA homologs present in the Actinobacteriophage Database (PhagesDB) (8). Sequences for Blino gp45, PCoral7 gp42, and Rebeelu gp30 are omitted as they are each 100% identical to CarolAnn gp44, Toast gp42, and Butters gp30, respectively. Sequence shading indicates conservation: white text on black background, 100% conserved; white text on dark gray background, 80% conserved; black text on light gray background, 60% conserved.

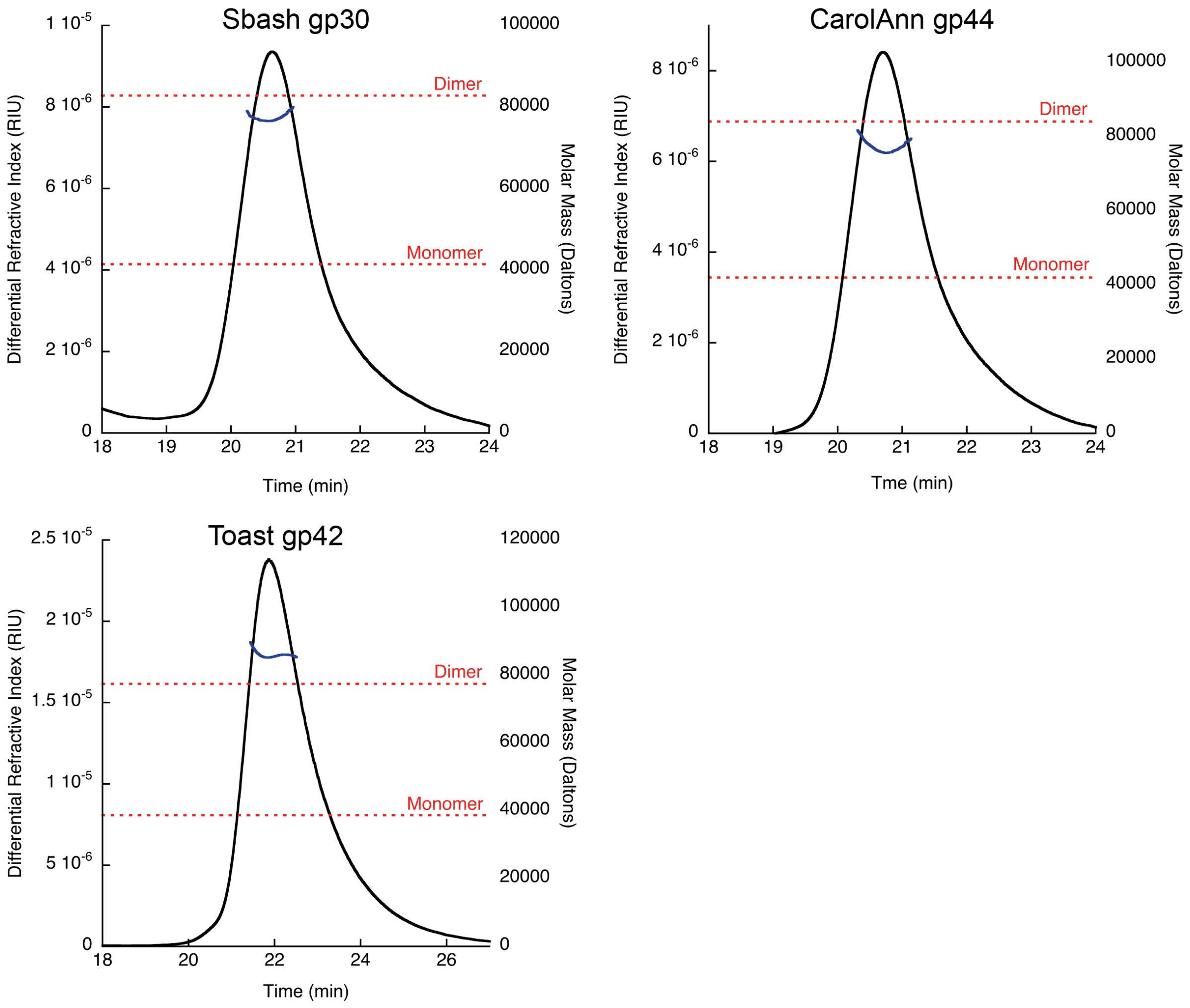


**Supplementary Fig. S10. SEC analysis of purified RexA homologs.** Black line denotes differential refractive index and blue line denotes measured mass across each peak. Dashed red lines indicate the predicted molecular weight of a monomer and dimer for each homolog.

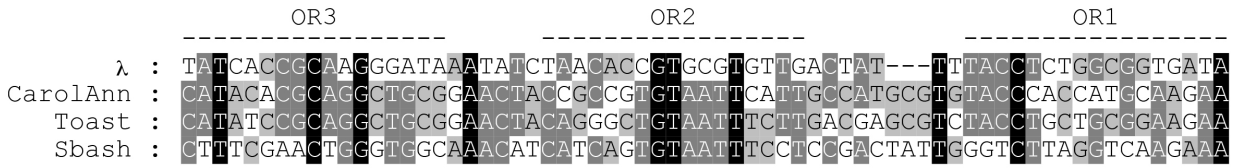


**Supplementary Fig. S11. Comparison of operator sequences used in binding experiments.** Alignment of the genomic region encompassing the operator sites $O_R1$, $O_R2$, and $O_R3$ in phages λ, CarolAnn, Toast, and Sbash. Sequence shading indicates conservation: white text on black background, 100% conserved; white text on dark gray background, 80% conserved; black text on light gray background, 60% conserved. DNA substrates containing λ OR1 and OR2 were used for EMSAs, limited proteolysis, and ESR experiments (**Figs. 4, 5, and 7**) while substrates containing OR1, OR2, and OR3 were only used for EMSAs with individual RexA homologs (**Fig. 7C**). See **Supplementary Table S2** for sequences of each individual substrate.

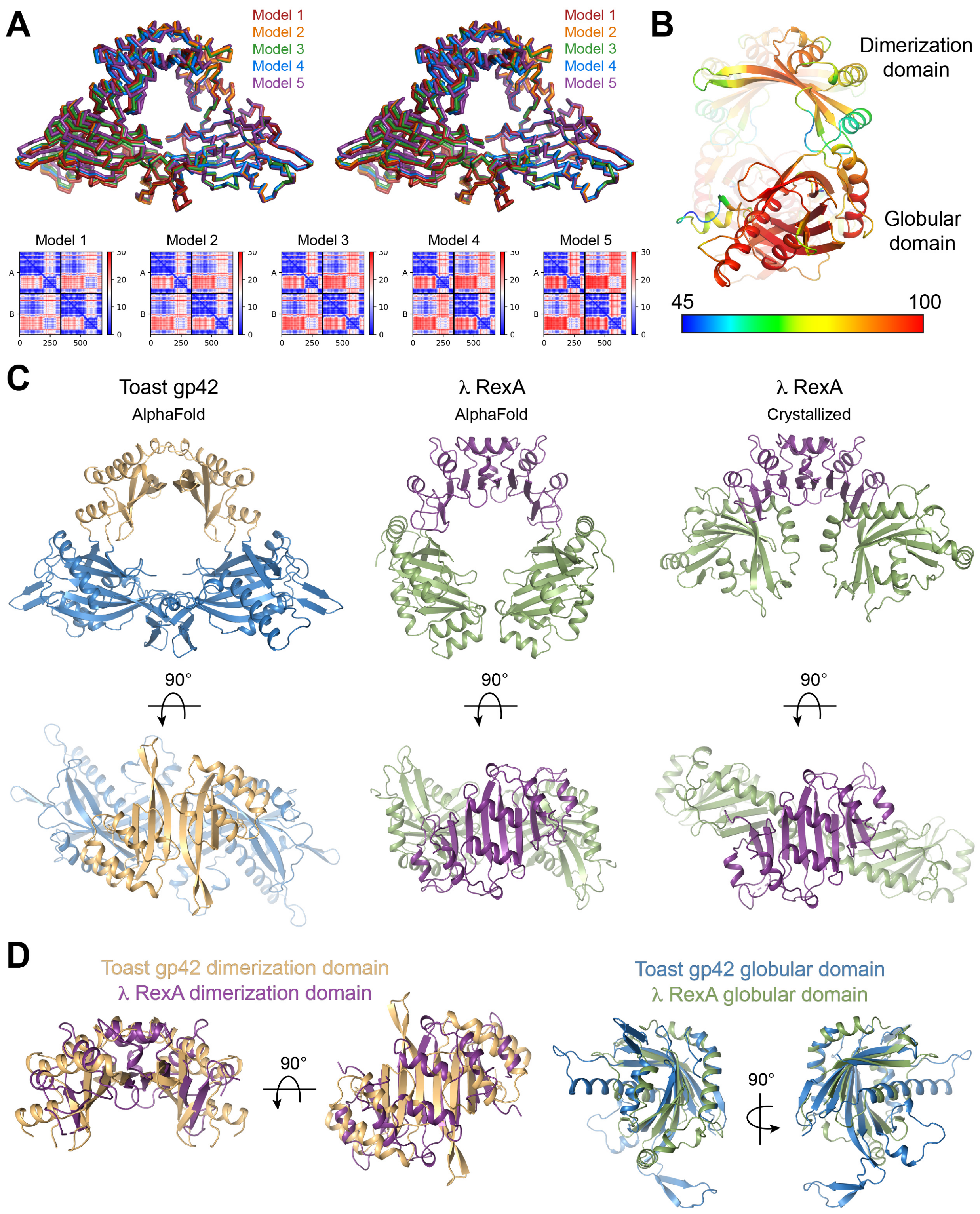


**Supplementary Fig. S12. AlphaFold model of Toast gp42 and comparison to RexA.** A. Superposition of top 5 Toast gp42 models generated with AlphaFold-Multimer (9) viewed in stereo. Predicted aligned error plots are shown below. B. Side view of Toast gp42 model colored according to the predicted local distance difference test (pLDDT) score (0-100), with values greater than 90 indicating high confidence and values below 50 indicating low confidence. Scale bar denotes per residue confidence coloring for pLDDT scores ranging from 45 to 100. C. Comparison of Toast gp42 AlphaFold model (left) with the RexA AlphaFold model (center, predicted open conformation) and crystallized RexA structure (right, closed conformation). Side and top views are shown for each. Coloring as follows: Toast gp42 dimerization domains, light orange; Toast gp42 globular domains, sky blue; RexA dimerization domains, violet purple; RexA globular domains, smudge. D. Superposition of the dimerization domains (left) and individual globular domains (right) from the Toast gp42 AlphaFold model and the crystallized RexA coordinates.
